# The formation of a fuzzy complex in the negative arm regulates the robustness of the circadian clock

**DOI:** 10.1101/2022.01.04.474980

**Authors:** Meaghan S. Jankowski, Daniel Griffith, Divya G. Shastry, Jacqueline F. Pelham, Garrett M. Ginell, Joshua Thomas, Pankaj Karande, Alex S. Holehouse, Jennifer M. Hurley

**Affiliations:** Department of Biological Sciences, Rensselaer Polytechnic Institute, Troy, NY, 12180, USA; Department of Biochemistry and Molecular Biophysics, Washington University School of Medicine, St. Louis, MO, 63110, USA; Department of Chemical and Biological Engineering, Rensselaer Polytechnic Institute, Troy, NY, 12180, USA; Center for Biotechnology and Interdisciplinary Sciences (CBIS), Rensselaer Polytechnic Institute, Troy, NY, 12180, USA; Center for Science and Engineering of Living Systems (CSELS), Washington University in St. Louis, St. Louis, MO, 63110, USA

**Keywords:** Intrinsic disorder, IDR, peptide microarray, LOCATE, electrostatics, molecular modeling, *Neurospora crassa*, FREQUENCY, Frequency-Interacting RNA Helicase, PERIOD, SLiM

## Abstract

The circadian clock times cellular processes to the day/night cycle via a Transcription-Translation negative Feedback Loop (TTFL). However, a mechanistic understanding of the negative arm in both the timing of the TTFL and its control of output is lacking. We posited that the formation of negative-arm protein complexes was fundamental to clock regulation stemming from the negative arm. Using a modified peptide microarray approach termed <u>L</u>inear m<u>o</u>tif dis<u>c</u>overy using r<u>at</u>ional d<u>e</u>sign (LOCATE), we characterized the interaction of the disordered negative-arm clock protein FREQUENCY to its partner protein FREQUENCY-Interacting RNA helicase. LOCATE identified a specific Short Linear Motif (SLiM) and interaction “hotspot” as well as positively charged “islands” that mediate electrostatic interactions, suggesting a model where negative arm proteins form a “fuzzy” complex essential for clock timing and robustness. Further analysis revealed that the positively charged islands were an evolutionarily conserved feature in higher eukaryotes and contributed to proper clock function.

## Introduction

Circadian clocks have evolved as an adaptive mechanism to anticipate daily environmental changes and are recognized as an important regulator of the cellular environment (Dunlap and Loros, 2017; Jankowski et al., 2020; Partch, 2020). The rhythms generated by these clocks are widely conserved amongst eukaryotes and their dysregulation leads to an increase in disease in humans (Dunlap and Loros, 2017; Evans and Davidson, 2013; Musiek and Holtzman, 2016; Shostak, 2017). Given its essential and conserved function across species, understanding the molecular mechanism that controls the timing of the circadian clock remains a fundamental question in biology. The underlying architecture of the circadian clock in higher eukaryotes (including fungi, insects, and mammals) is a molecular oscillator composed of a Transcription-Translation negative Feedback Loop (TTFL), made up of a positive transcriptional-activating protein complex and a negative repressing protein complex (Figure 1A) (Darlington et al., 1998; Dunlap and Loros, 2017; Gekakis et al., 1998; Hurley et al., 2016). The positive arm of the clock plays a role in setting off a cascade of transcriptional activation that modulates the daily expression of up to 80% of transcripts at the organismal level, while the negative arm represses positive arm activity (Dunlap and Loros, 2017; Hurley et al., 2014; Miller et al., 2007; Mure et al., 2018; Tataroglu and Emery, 2014). Despite an understanding of the emergent properties that circadian clocks encode on a cellular scale, questions remain regarding the fundamental biophysical mechanisms of feedback and circadian post-transcriptional regulation (Collins et al., 2021; Hurley et al., 2018; Kojima et al., 2011; Mauvoisin et al., 2014; Mauvoisin and Gachon, 2020; Partch, 2020; Reddy et al., 2006).

**Figure 1.**
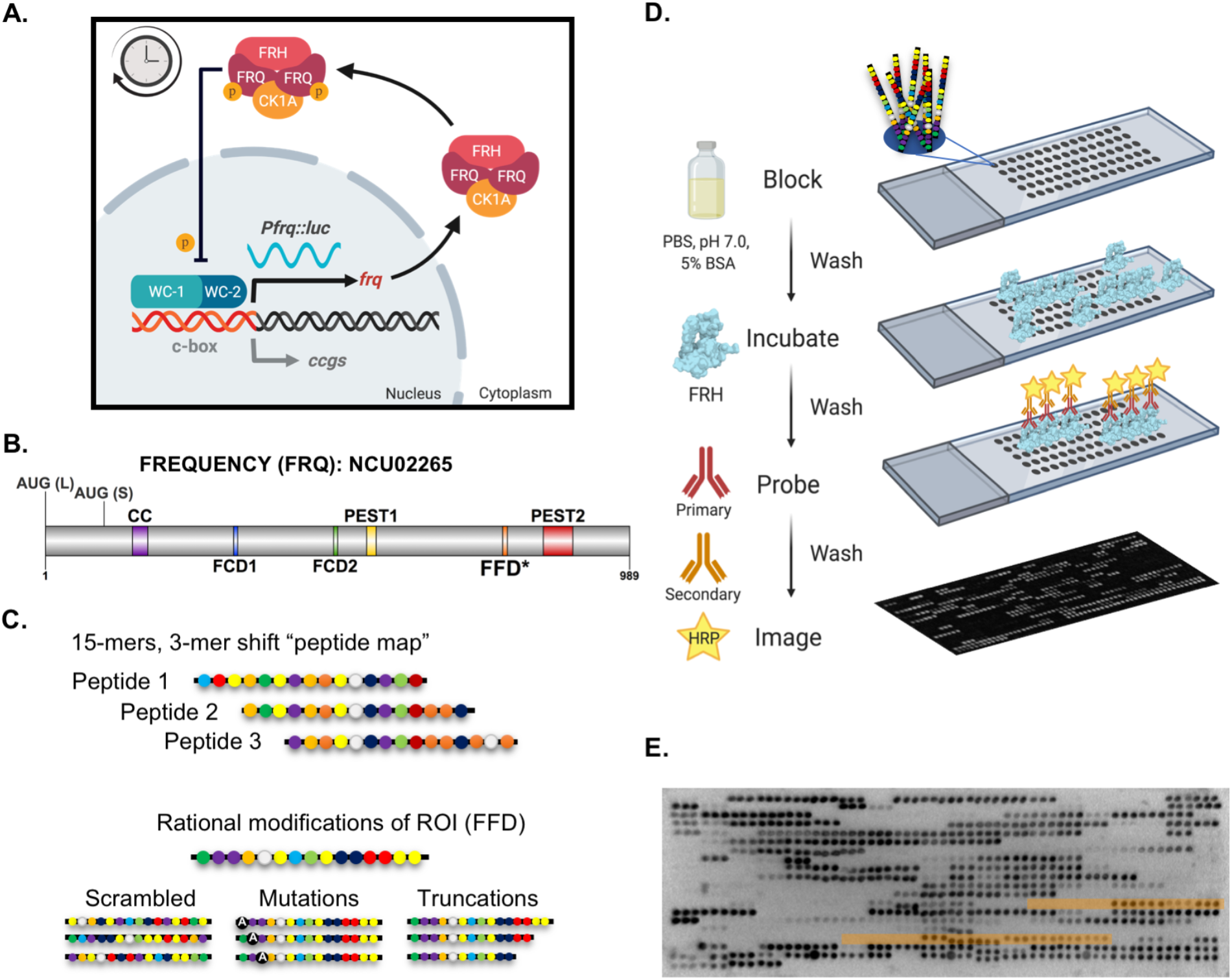
LOCATE maps FRQ/FRH interactions at the single residue level. (A) Schematic of the molecular clock in *Neurospora crassa.* The transcriptional activators WC-1 and WC-2 activate *frq* transcription. Once translated, FRQ binds CK-1A and FRH leading to repression of WC-1/WC-2 activity at the *frq* promoter, presumably by phosphorylation, closing the feedback loop. Robust oscillation of the clock is represented by the continuous oscillation of luciferase expressed from the *frq* promoter in our reporter strain. FRQ = FREQUENCY, CK1a = CASEIN KINASE 1a, FRH = FRQ-INTERACTING RNA HELICASE, Pfrq = frq promoter, luc = luciferase, c-box = clock box, ccgs = clock-controlled genes, p = phosphorylation. (B) FRQ protein topology with known binding domains highlighted, AUG (L) = Long FRQ start codon, AUG (S) = Short FRQ start codon (Garceau et al., 1997), CC = Coiled-coil region (Cheng et al., 2001), FCD1/2 = FRQ-CK-1A interaction Domain 1/2 (Querfurth et al., 2011), PEST 1/2 = Proline-Glutamic Acid-Serine-Threonine rich region 1/2 (Görl et al., 2001), FFD = FRQ-FRH interacting Domain, * = region as defined by Guo et al. 2010). Plotted using DOG (v2.0) (Ren et al., 2009). (C) The design of the FRQ LOCATE peptide library based on a 15-mer “peptide map” scan, shifting by 3 a.a., through the primary sequence of FRQ. Rational modifications were designed for Regions of Interest (ROI) using scrambled sequences, mutations, and truncations. (D) The experimental workflow for the LOCATE method; blocking, incubating with FRH, probing using an FRH-specific antibody, and chemiluminescent detection. (E) Example of a FRQ microarray challenged with ∼10 nM FRH and probed with anti-His antibodies. Image inverted for density analysis of FRH binding. Regions highlighted in orange are peptides including the FFD region (3-mer shift vs. 1-mer shift). Related to Supplemental Figure 1.

Clues to feedback and post-transcriptional regulatory mechanisms have come from the extensive, ontologically diverse, macromolecular protein complexes that coalesce around the proteins that comprise the negative arm of the clock (reviewed in Mosier and Hurley, 2021)(Baker et al., 2009; Duong et al., 2011; Oyama et al., 2019; Pelham et al., 2021). Negative-arm centered interactomes are likely enabled by their enrichment for Intrinsically Disordered Regions (IDRs) in the negative arm proteins which are characterized by flexible polypeptide backbones, a lack of a fixed tertiary structure, and the formation of dynamic protein-protein interactions (Partch et al., 2005; Hurley et al., 2013; van der Lee et al., 2014; Wright and Dyson, 2015; Pelham et al., 2020; Uversky et al., 2000). Proteins with IDRs frequently engage in multivalent intermolecular interactions, generally functioning as hubs for cellular information processing (Cermakova et al., 2021; Cortese et al., 2008; Cumberworth et al., 2013; Haynes et al., 2006; Martin and Holehouse, 2020; Wright and Dyson, 2015). While IDR-based protein-protein interactions may provide insights into potential regulatory mechanisms in the clock, technical limitations hamper biophysical studies of the interactions that occur across IDRs in a large protein, constraining the investigation of this potentially important circadian regulatory mechanism (Mendoza and Vachet, 2009; Schneider et al., 2019; Suskiewicz et al., 2011; Uemura et al., 2018).

Currently, our understanding of the interactions within negative-arm clock protein complexes stems mainly from genetic studies that assume interaction regions occur primarily within conserved domains (Akashi et al., 2014; Guo et al., 2010; Kim et al., 2007; Querfurth et al., 2011). However, proteins with IDRs commonly engage in molecular interactions via short (5-15 residue), degenerate regions termed <u>S</u>hort <u>Li</u>near <u>M</u>otifs (SLiMs) (Bugge et al., 2020; Davey et al., 2012; Edwards and Palopoli, 2015; Gouw et al., 2017; Obenauer et al., 2003; Ren et al., 2008; Sigrist et al., 2002; van der Lee et al., 2014). Taking into account the large IDRs found in many clock proteins, we theorized that hidden and/or degenerate SLiMs that lack conventional signatures of conservation play key roles in dictating clock-associated molecular interactions, and thereby feedback and regulatory output mechanisms (Pelham et al., 2020). Synthetic peptide microarrays have been used to successfully characterize SLiMs that occur within a single IDR, by using a microarray of printed peptides to measure the binding of a protein, or structured domain, to an IDR (Blikstad and Ivarsson, 2015; Harnoš et al., 2018; Kumar et al., 2017). Given the success of this approach, we hypothesized that this “divide-and-conquer” strategy could be extended to investigate IDRs in the clock negative-arm proteins to grant insight into negative-arm protein-protein interactions and, by extension, the mechanisms of feedback and output regulation (Katz et al., 2011; Sethi et al., 2012).

Here we report the application of synthetic peptide microarrays for the study of protein-protein interactions involving proteins enriched in IDRs. Our approach, termed <u>L</u>inear m<u>o</u>tif dis<u>c</u>overy using r<u>at</u>ional d<u>e</u>sign (LOCATE), used rationally designed synthetic peptides printed in a microarray format to perform a high-throughput screen of a large disordered protein to characterize interaction domains (Shastry and Karande, 2019). Using the largely disordered negative arm clock protein FREQUENCY (FRQ) from *Neurospora crassa* (*N. crassa*) as a model, LOCATE characterized the binding relationship between FRQ and its partner protein FRQ-interacting RNA Helicase (FRH), revealing that beyond the FRH-specific SLiM, FRH bound to multiple positively charged “islands” within FRQ (Cheng et al., 2005; Hurley et al., 2013). This multivalent interaction aligned with a model of a “fuzzy” FRQ/FRH complex that could affect the availability of other predicted FRQ SLiMs for downstream interactions. We found that positively charged islands were a conserved feature in the negative arm of higher eukaryotes (e.g., the functionally orthologous PER protein family), suggesting that distributed electrostatic interactions, and perhaps fuzzy complexes, are an evolutionarily conserved feature of negative arm proteins. LOCATE further identified “hotspot” residues that facilitated binding of the FRH-specific SLiM that, once mutated, allowed us to demonstrate the FRQ/FRH multivalent interaction is essential for clock robustness rather than repression. Our results point to a mechanism in which positively-charged residues can generate an electrostatically favorable context to enable distinct modes of interaction that mediate recognition and binding between clock proteins.

## Results

### LOCATE is an unbiased method to identify established binding motifs and novel interaction regions

To characterize binding between negative-arm clock protein IDRs and their interactors, we adapted a peptide microarray-based method we designated LOCATE (Shastry and Karande, 2019). Employing a peptide library based on the primary sequence of a largely disordered target protein, the LOCATE method can identify SLiMs and other sequence elements that contribute to binding (Davey et al., 2012; Harnoš et al., 2018; Katz et al., 2011; Shastry and Karande, 2019; Van Roey et al., 2014). We applied LOCATE to the *N. crassa* negative arm protein FREQUENCY (FRQ, NCU02265), as a model system to test this approach. With the exception of several conserved interaction domains, FRQ is largely intrinsically disordered, binds many different partners, and contains many putative SLiMs (Figure 1B, Supplemental Figures 1A and 2D) (Baker et al., 2009; Garceau et al., 1997; Guo et al., 2010; Hurley et al., 2013; Pelham et al., 2021; Querfurth et al., 2011).

We constructed a 15-mer “peptide map” of sequentially overlapping peptides that scanned through the primary sequence of FRQ, shifting by 3 a.a. each time (Figure 1C). Beyond a full peptide map, specific regions of interest were investigated by rationally designed peptides to test binding specificity (e.g., scrambles, single-point mutations, and truncations) (Figure 1C) (Cheng et al., 2001; Görl et al., 2001; Guo et al., 2010; Querfurth et al., 2011; Shastry and Karande, 2019). Peptide libraries were synthesized using standard Fluorenylmethyloxycarbonyl (Fmoc) chemistry and printed in peptide microarray format, as previously described (Shastry and Karande, 2019). Peptides were synthesized from C- to N-terminus on a solid surface and, other than the peptide taken from the very N-terminus of FRQ, all peptides were N-terminally acetylated to maintain a more native overall charge. Following a solubilization step, peptides were spotted in triplicate onto a microarray slide.

To validate the LOCATE approach, we examined the binding between the FRQ library and the well-studied interactor of FRQ, FREQUENCY-Interacting RNA Helicase (FRH, NCU03363). We incubated the printed FRQ LOCATE microarray with *E. coli* expressed FRH (a.a. 110-1106, with a 6x His-tag) (Figure 1D and Supplemental Figure 1B) (Conrad et al., 2016). Using an FRH-polyclonal antibody (Shi et al., 2010), as well as a monoclonal anti-His antibody, FRH binding to each FRQ-based peptide was assessed using chemiluminescent imaging. As expected, there were single peptides (triplicate dots) that showed stochastic FRH-binding behavior (Figure 1E), which can be due to synthesis issues or other artifacts that lead to false positive binding with synthetic peptides (Blikstad and Ivarsson, 2015). Discounting these spuriously-bound peptides, we replicated the previous genetically-identified FRQ-FRH domain (FFD, a.a. 774-782) as a significant region of interaction (Figure 1E, areas highlighted in orange) (Guo et al., 2010). We also found many instances where sequential peptides showed increased and then decreased signal as a set of amino acid sequences entered and exited the 15 a.a. sliding window, suggesting FRQ/FRH interaction was more complex and involved regions beyond the FFD (Figure 1E) (Cheng et al., 2005; Guo et al., 2010).

### FRQ binds FRH via distributed islands of positively charged residues

We first investigated the composition of the top-binding peptides relative to the FRQ LOCATE 3-mer shift peptide map to characterize the FRQ/FRH binding regions beyond the FFD. We found that the top 10% of FRH-binding peptides were enriched in basic residues, predominantly arginine, and significantly depleted in acidic residues (Figures 2A and B). We also noted that FRH binding across the FRQ peptide-mapping library was skewed towards more positively charged peptides (Figure 2C). This correlated with two previously-solved FRH crystal structures that show the surface electrostatic potential of FRH is mainly negative, with the exception of the positive RNA-binding groove and a few other isolated domains (e.g. the KOW binding domain) (Figure 2D) (Conrad et al., 2016; Morales et al., 2018).

**Figure 2.**
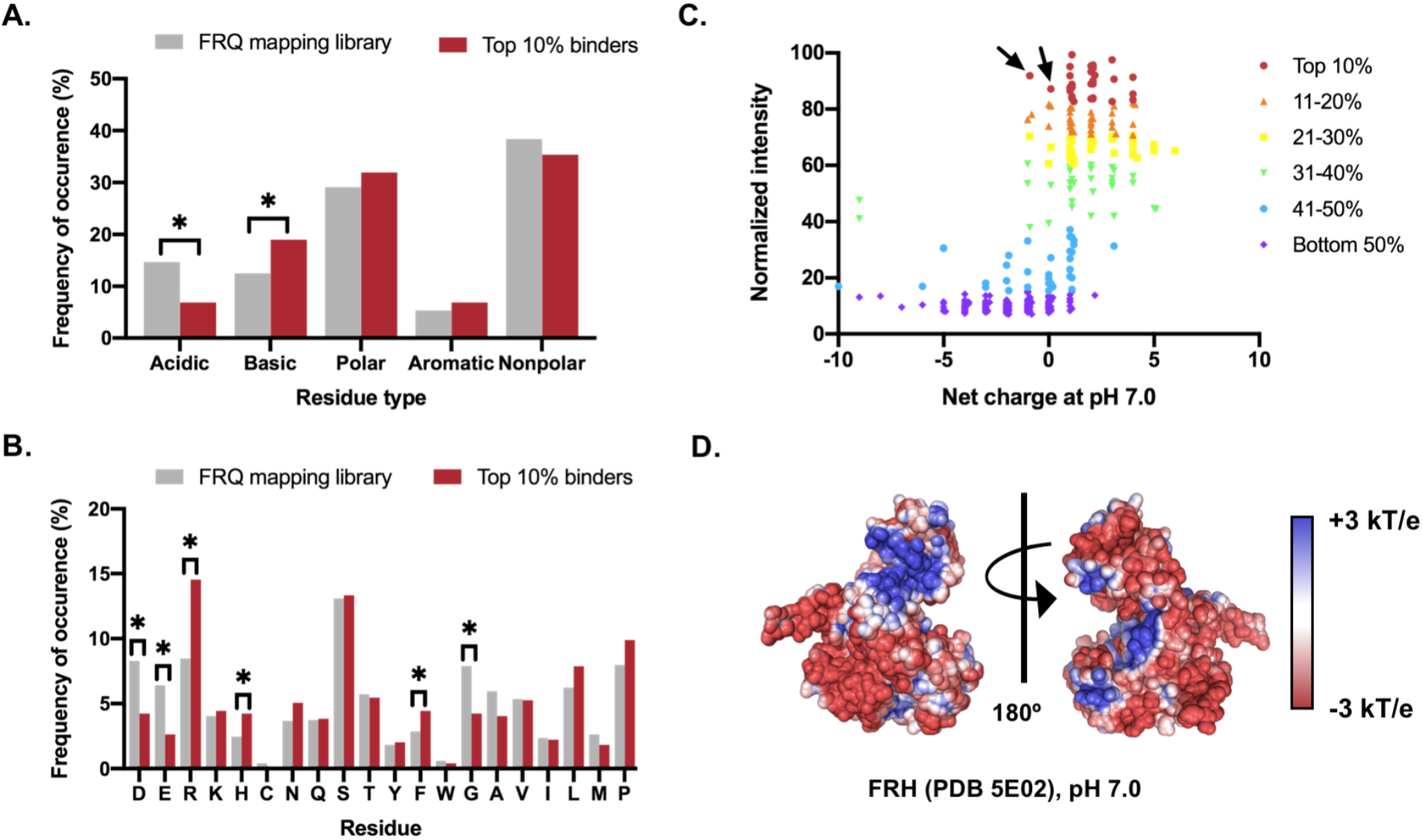
Charge residues contribute to the interaction between FRQ and FRH. (A) Comparison of the amino acid types in the overall 3-mer shift mapping library and the amino acid types in the top 10% of binding peptides (two-tailed z test for population proportions; * = p<0.05). Acidic = D and E; basic = R and K; polar = H, C, N, Q, and S; aromatic = Y, F, and W; and nonpolar = G, A, V, I, L, M, and P. (B) Comparison of the amino acids in the overall 3-mer shift mapping library and the amino acids in the top 10% of binding peptides (two-tailed z test for population proportions; * = p<0.05). Statistical significance of frequency changes of residues C, Y, W and M was not determined due to low residue occurrence in the sample population. (C) Ranking of peptides by normalized binding-intensity and estimated net charge at pH 7.0. Peptides colored by their binding decile as indicated. Arrows denote two peptides containing the FFD with an overall neutral charge. Note (A) to (C) was based on average normalized intensities from anti-His and anti-FRH experiments using ∼10 nM FRH. (D) The calculated surface electrostatic potential map for FRH. Blue denotes positive charge while red denotes negative charge, ranging from +3 kT/e to −3 kT/e.

To evaluate the role of charge in FRQ/FRH binding, we next compared FRH binding regions on FRQ with the linear distribution of the Net-Charge Per Residue (NCPR) across the sequence of FRQ. FRH binding correlated strongly with regions of high positive NCPR on FRQ (Figure 3A). This analysis showed that positively and negatively charged residues were interspersed on FRQ in clusters, or islands (Figure 3A). The relative position of oppositely charged residues (charge patterning) has emerged as an important sequence feature for highly disordered proteins (Cohan et al., 2021; Das and Pappu, 2013; Sawle and Ghosh, 2015; Sherry et al., 2017). While charge patterning can be quantified using the parameter κ (kappa) and sequence charge decoration (SCD), we reasoned that as FRH is predominantly negatively charged, the positively charged islands within FRQ would be the most relevant to binding (Cohan et al., 2021; Das and Pappu, 2013; Sawle and Ghosh, 2015; Sherry et al., 2017). We therefore focused on quantifying the extent of positive charge clusters using the established Inverse-Weighted Distance (IWD) measure, which takes the composition and sequence distribution of any single residue type into account (Eq. 2 in methods) (Holehouse et al., 2021). By taking the IWD value for positive residues within the FRQ sequence and comparing it to the IWD values calculated from 10,000 randomly shuffled FRQ sequences, we found that positively charged residues in FRQ were significantly more clustered together than expected by random chance (Figure 3B) (Cohan et al., 2021; Holehouse et al., 2021; Martin and Holehouse, 2020; Schueler-Furman and Baker, 2003). This clustering was also seen for negative residues, but not aromatic residues, within FRQ (Supplemental Figure 2A and B).

**Figure 3.**
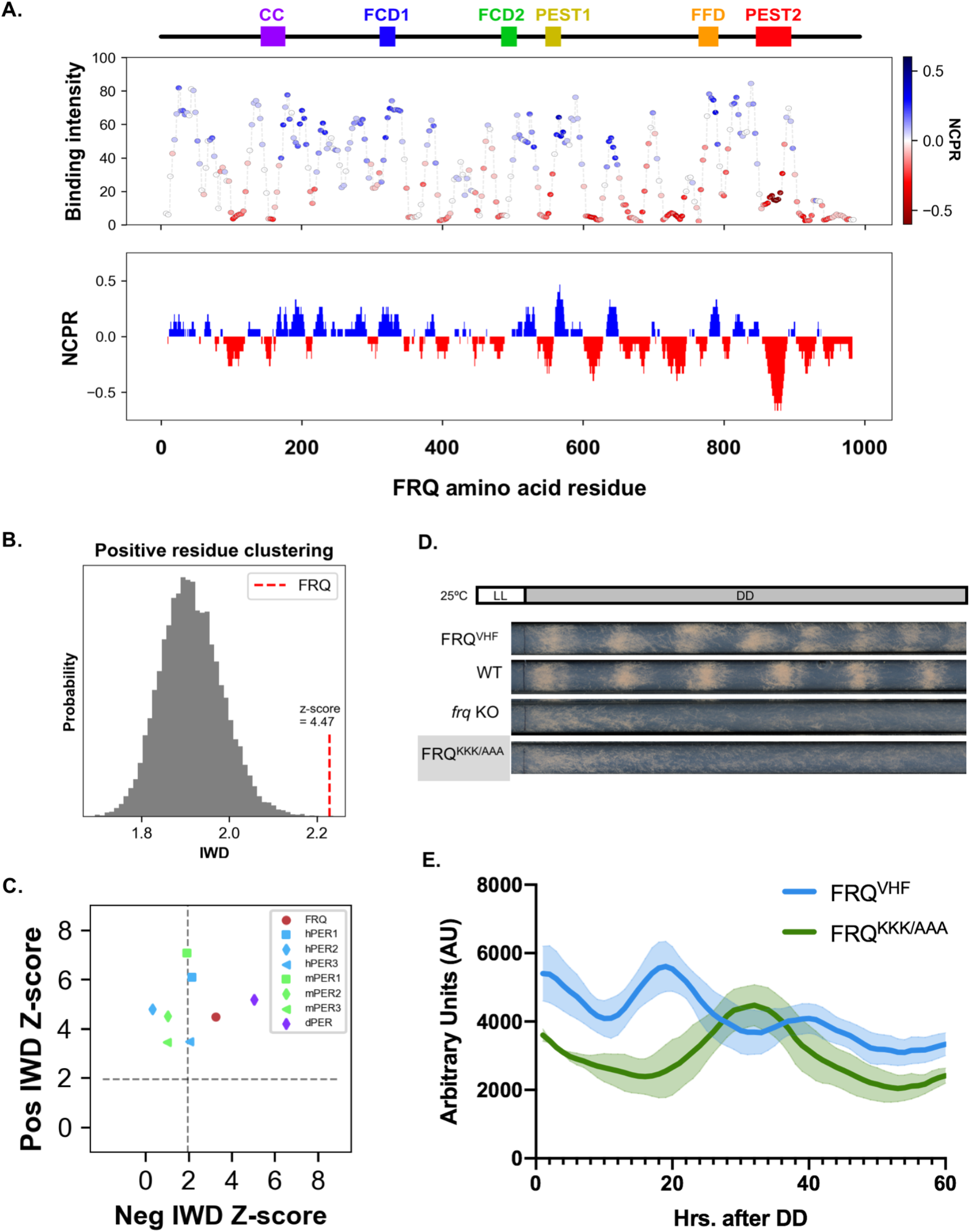
FRQ/FRH binding occurs within positively charged islands. (A) Normalized average FRH-binding intensity (∼10 nM FRH, anti-His), with peptides color coded by net charge per residue (NCPR) (blue = positive NCPR and red = negative NCPR), and local NCPR (using a 15 residue sliding window) plotted against the sequence and known domains of FRQ (abbreviations as in Figure 1). (B) The Inverse Weighted Distance (IWD) parameter to measure positive residue clustering in FRQ (red) compared to the IWD null distribution (gray) based on 10,000 FRQ sequence shuffles. (C) The normalized Z-score for the calculated IWD parameter for positive and negative residues for FRQ (red circle) compared to the IWD Z-scores of PER orthologs (see legend). The dashed lines denote significance, with values greater than this line being significant at p<0.05. (C) Representative race tubes of FRQ^VHF^, Wild-type FRQ (WT), *frq* KO and FRQ^KKK/AAA^ strains grown in DD. (D) Average and standard deviation values of N=3 FRQ^VHF^ (tagged wild-type) vs. FRQ^KKK/AAA^ strains with a luciferase reporter of *frq* promoter activity (*Pfrq(c-box)::luc*) background grown in DD in a 96 well format in the presence of luciferin. Related to Supplemental Figures 2 and 3.

To determine if the extensive phosphorylation of FRQ over the circadian day could modulate these charge islands, we calculated how charge islands were changed by temporal phosphorylation (Baker et al., 2009). We found that positive charge clusters were, in general, maintained, even when all FRQ’s detected phosphosites were considered negatively charged (Supplemental Figure 2C). This lead us to hypothesize that charge islands aligned with the location of predicted SLiMs for other verified FRQ-binding proteins (Pelham et al., 2021). Our analysis showed that FRQ SLiMs tended to occur near negatively charged islands, suggesting a relationship between FRQ/FRH binding and the formation of FRQ macromolecular complexes (Supplemental Figures 2C and D) (Pelham et al., 2021).

Finally, we analyzed the IWD for different orthologs of FRQ to investigate if charge islands are a conserved evolutionary feature of the circadian negative arm proteins. We transformed the IWD into Z-scores using a null background of 10,000 randomly shuffled sequences of the same composition to allow a more direct comparison between orthologs (Cohan et al., 2021; Martin and Holehouse, 2020). Like FRQ in *N. crassa*, most FRQ orthologs contained significant positive clusters (Supplemental Figure 3A). The functional PERIOD (PER) ortholog proteins in *D. melanogaster*, *M. musculus*, and *H. sapiens* also demonstrated significant positive charge clustering (Figure 3C). In contrast, negative charge clustering was significant in only a few orthologs, suggesting that positive charge clustering may be a conserved molecular feature amongst disordered negative arm clock proteins in higher eukaryotes (Zarin et al., 2019).

### Charged islands in FRQ support the proper timing of the clock feedback loop

To investigate the importance of FRH binding via positively charged islands on the function of the circadian clock, we analyzed the effect of the mutation of residues in a positively charged island (KKK, a.a. 315-317) adjacent to the genetically identified CK-1A interaction site known as FCD-1 (Querfurth et al., 2011). FCD-1 is proposed to interact through a hydrophobic helical interface defined by residues 319-326, and the KKK residues fall just outside of the canonical FCD-1 motif (Querfurth et al., 2011). Given the observation that positively charged islands are distributed across FRQ, we hypothesized that modulating positively charged residues in an island that flanks a known binding motif might impact function, confirming the importance of the positively charged islands.

In order to test our hypothesis, we took advantage of tools in *N. crassa* that allow for the analysis of clock function *in vivo*. We targeted VHF-tagged (V5, 10-His, 3-Flag) alleles of FRQ to the cyclosporin (*csr-1*) locus of an *frq* KO strain that had the banding mutation and a luciferase reporter fused to the *frq* promoter (*pfrq:luc*) in its genetic background, creating the wildtype (FRQ^VHF^) and KKK315AAA (FRQ^KKK/AAA^) strains (Bardiya and Shiu, 2007; Belden et al., 2007; Emerson et al., 2015; Gooch et al., 2014; Sargent and Woodward, 1969). Banding mutants (bd+) of *N. crassa* are known to lay down one conidial “band” per clock cycle when grown on a layer of agar-based media within a hollow glass “race tube”, allowing us to test the effect of the KKK-AAA mutation on the overt clock (Sargent and Woodward, 1969). Analysis of the formation of conidial bands in the FRQ^KKK/AAA^ strain demonstrated that, while the FRQ^VHF^ strain maintained a clock, the FRQ^KKK/AAA^ strain had no overt clock rhythms (Figure 3D and Supplemental Figure 3B).

Banding is a measurement of clock output but is not always representative of the activity of the core clock. Therefore, we next assayed *frq* promoter activity using the luciferase reporter present in the strains as a proxy for the TTFL, as the activation of the *frq* promoter leads to the transcription/translation of luciferase (Gooch et al., 2014). Compared to the FRQ^VHF^ strain, which showed a typical ∼21 hr. oscillation in *frq* promoter activity as reported by luminescence, the *frq* promoter activity for the FRQ^KKK/AAA^ strain displayed a non-circadian oscillation (approx. 48 hrs) (Figure 3E and Supplemental Figure 3C). The loss of clock function due to the KKK to AAA mutation supports a model in which positively charged residues can potentiate or steer conserved hydrophobic binding motifs on FRQ by generating an appropriate sequence context. To further explore this idea, we turned to a second known binding motif in FRQ, the FFD.

### Hotspot residues C-terminal of the FFD are critical for FRH binding in vitro

While the overall binding behavior of FRH to FRQ appeared to be related to electrostatics, LOCATE highlighted FRH binding to several FRQ peptides that had a net neutral charge (see arrows in Figure 2C) (Guo et al., 2010). These peptides mapped to the FFD, a previously characterized FRH-interaction region (a.a. 774-782) (Figures 4A and B). Unlike other FRH-binding peptides identified by LOCATE, the FFD is composed of multiple hydrophobic residues at its center and flanked by polar or charged residues (Figure 4A). Amongst FRQ homologs, the exact FFD motif, or SLiM, is not conserved, but similar physicochemical residues are retained at several positions, suggesting an evolutionary chemical signature (Figure 4A and Supplemental Figure 4A). When we analyzed the binding of FRH to scrambled FFD peptides that maintained the same amino acid composition and neutral net charge, we found scrambles with diminished binding (Supplemental Figure 4B). We interpret this to mean the net charge of the FFD motif is not the only determinant of binding. In contrast, a comparable analysis of the electrostatically-driven interaction near the FCD-1 region (described above, peptide MTDKE**KKK**LVVRRLE, predicted net charge +3) showed minimal changes in binding intensity upon shuffling (Supplemental Figure 4C). These analyses distinguish the 15-residue FFD peptide from other FRH-binding peptides as a region that is not solely dependent upon electrostatic interactions between FRQ and FRH and may have some sequence-specificity.

**Figure 4.**
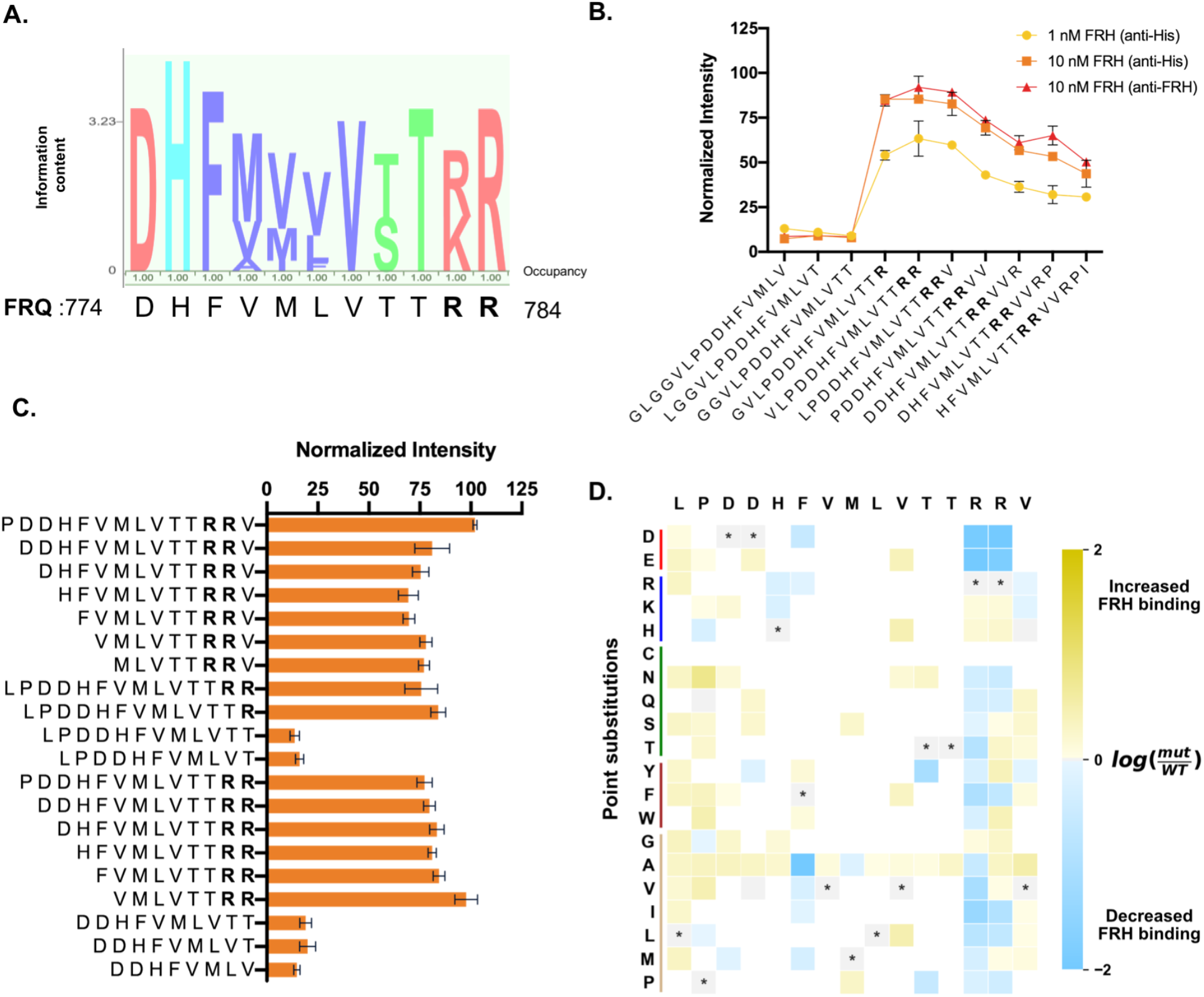
A LOCATE-identified novel conserved RR region is essential for FRQ/FRH interaction. (A) Weighted observed residues at each position for the FFD region of FRQ over 10 FRQ homologues, with *N. crassa* FRQ residues below. Color denotes residue type (pink is charged, blue is histidine or tyrosine, purple is small/hydrophobic, and green is a hydroxyl/amine following ClustalX scheme). Occupancy refers to the percent the position was filled. (B) The average normalized intensity of FRH binding to the stated FRQ-based peptides from Library I. (C) Average normalized microarray intensity of FRH-binding for each FFD-based peptide truncation tested. (D) Point substitution analysis of the FFD SLiM with the original peptide listed along the top and the tested amino acid point substitution along the left-hand side. Grayed boxes with stars denote the wild-type residue at that position, while yellow denotes an increase in FRH associated with a peptide that has a given substitution at that position, while blue denotes a decrease in FRH binding intensity. Note that the scale is log_2_ of mutant peptide (mut)/wild type peptide (WT). White boxes correspond to amino acids substitutions that were not tested. Unless otherwise noted, results in this figure are based on Library II peptides, incubated with ∼100 nM FRH and visualized with anti-His. Error bars denote standard deviation. Related to Supplemental Figure 4.

SLiM-based protein interactions are often driven by specific residues within the motif, termed hotspots (Bogan and Thorn, 1998). To identify single or multiple hotspot residues within the FFD SLiM, we scrutinized the FFD motif in our LOCATE analysis. We noted a sudden increase in binding when two arginines C-terminal of the canonical FFD motif entered the peptide window (a.a. 783-784, Figure 4B). To determine if these arginines were hotspot residues, we made rational mutations (truncations and point mutations) of a peptide that included both the FFD and the double arginines. We found that the arginines were critical for FRQ/FRH binding, suggesting that the previous definition of the FFD motif (a.a. 774-782) was incomplete (Figure 4C and D, Supplemental File 1). This result is in keeping with the over-representation of arginine (R) in the top 10% of FRH-binding peptides and is consistent with the finding that arginine is the second most common amino acid in protein interaction regions (Figure 2B) (Bogan and Thorn, 1998; Mendoza and Vachet, 2009).

### The FFD arginine hotspot residues are essential for FRQ/FRH binding in vivo

To verify the importance of the LOCATE-identified RR hotspot residues *in vivo*, we again targeted VHF-tagged (V5, 10-His, 3-Flag) alleles of FRQ to the cyclosporin (*csr-1*) locus of an *frq* KO strain (Figure 5A) (Bardiya and Shiu, 2007; Emerson et al., 2015). We replicated a previously published alanine substitution of part of the FFD region before the arginine residues, known to break FRQ/FRH binding (FRQ^FFD2^, VMLVTT-AAAAAA), and created double arginine to double alanine (RR783AA, FRQ^RR/AA^) and double arginine to double histidine (RR783HH, FRQ^RR/HH^) mutant strains (Figure 5A) (Guo et al., 2010). Relative to the FRQ^VHF^ strain, low levels of FRQ^FFD2^ were present, similar to what was observed previously (Figure 5B) (Guo et al., 2010). We tested the ability of these FRQ isoforms to interact with FRH *in vivo* using co-immunoprecipitation (Figure 5B and Supplemental Figure 5A). While FRQ^VHF^ pulled down ample FRH, as did FRQ^RR/HH^, as predicted by LOCATE and previous publications, both FRQ^FFD2^ and FRQ^RR/AA^ were unable to co-immunoprecipitate FRH (Figure 5B and Supplemental Figure 5A) (Guo et al., 2010).

**Figure 5.**
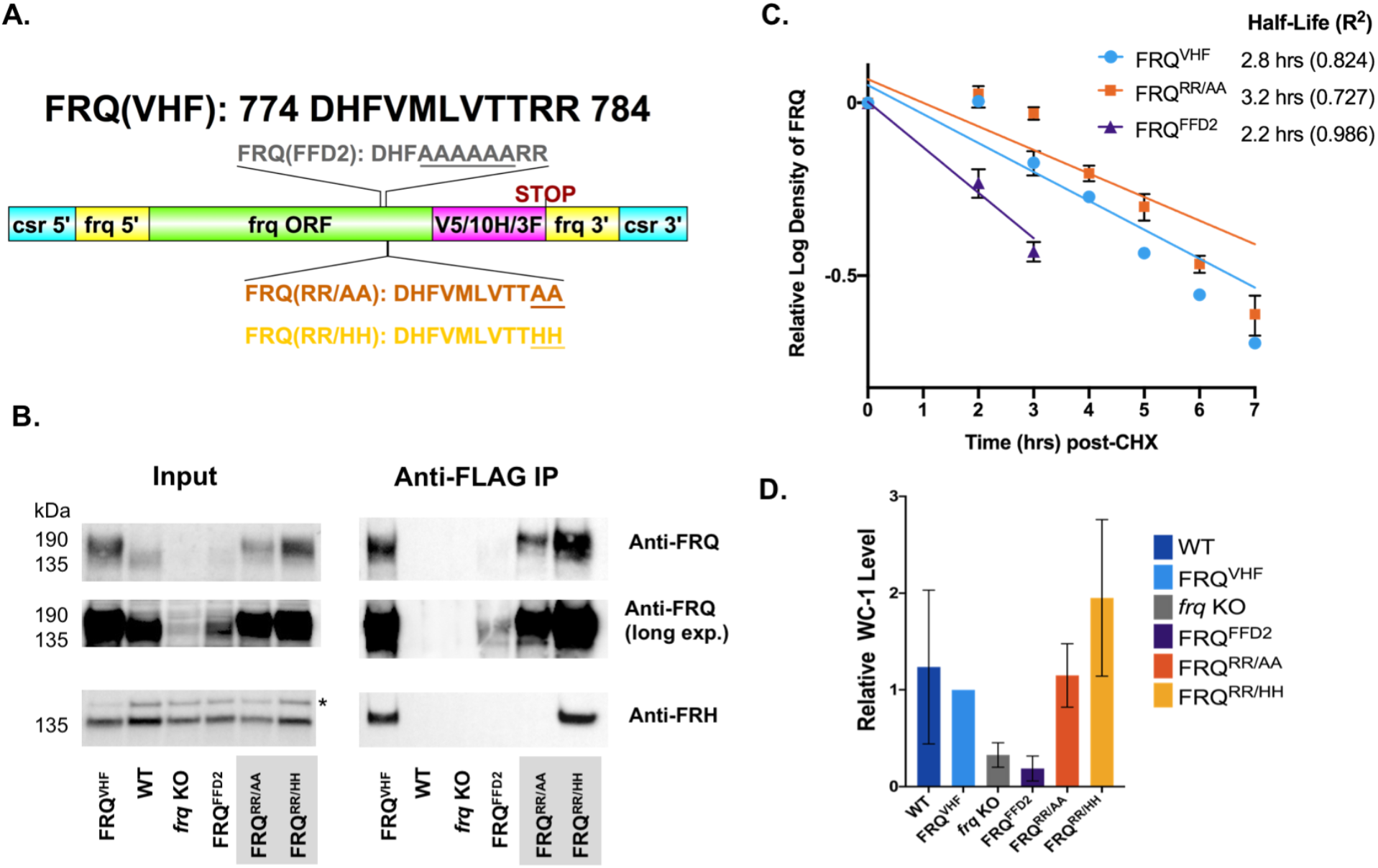
*In vivo* mutation of the LOCATE-identified RR residues leads to a loss of FRQ-FRH interaction but no loss in FRQ stability. (A) Schematic of the genetic mutations made within the FRQ-FFD region. Plotted using DOG (v2.0) (Ren et al., 2009). (B) Western blot of the anti-Flag co-immunoprecipitation of the described VHF-tagged FRQ strains. (C) Semi-quantitative FRQ half-life analysis based on a cycloheximide assay assuming one-phase decay. (D) Densitometry analysis of WC-1 lysate levels in the described VHF-tagged FRQ strains. Error bars denote standard deviation. Related to Supplemental Figures 5 and 6.

We next investigated how the double arginine residues might contribute to the FFD motif’s binding affinity. Shuffle mutants did not support a sequence-specific binding interface, as some shuffled variants that reposition the arginine residues within the motif bound with wildtype-like affinity (Supplementary Figure 4B). To gain mechanistic insight into how the double arginine region could affect the binding of FRQ to FRH, we performed all-atom simulations of a 50 a.a. section of FRQ that centered on the FFD region (Supplemental Figures 5B-E, Supplemental Movie 1). Simulations of wildtype (WT), RR783AA, RR783HH with a neutral histidine, and RR783HH with a positively charged histidine, were performed. The RR to AA and HH variants had a limited impact on the structural ensemble of the region outside of the specific position where the mutations occur (Supplemental Figure 5B,C,D). Taken naively, these simulations imply that the loss of arginine residues does not lead to a major change in the overall conformational behaviour of the canonical FFD. As such, our data seem to be best described by a model in which the arginine residues contribute an electrostatic component to the specific binding noted in the earlier region of the FFD. As the local context of a motif, such as charged residues, has been noted as important for the specificity of interactions, our results here further support a model in which the double arginine hotspot mediates essential electrostatic interactions that likely work in combination with the hydrophobic residues to facilitate FRH binding to the FFD region (Bugge et al., 2020; Ivarsson and Jemth, 2019).

Overall, our results implied that FRQ binding to FRH was driven by distributed, multivalent binding sites in which both charged and hydrophobic residues can contribute in ways that depend on both the sequence order and the sequence chemistry. The functional consequences of deleting the hydrophobic residues in FFD are well-established (Guo et al., 2010). However, we wondered if disrupting FFD-mediated interaction by mutating the arginine hotspot residues would have the same phenotypic effect, or if distinct outcomes can be achieved by altering IDR chemistry in orthogonal ways.

### Mutation of the FFD hotspot residues stabilizes FRQ in vivo

A reduction in FRQ/FRH interaction, as seen in the FRQ^FFD2^ and FRQ^RR/AA^ mutants above, has been correlated with a loss of FRQ stability (Cheng et al., 2005; Guo et al., 2010; Hurley et al., 2013). This phenomenon has been attributed to FRH’s function as a Nanny protein, preventing constitutive degradation (‘degradation by default’) via an E3 ubiquitin ligase-independent protein degradation pathway (He et al., 2003; Hurley et al., 2013; Suskiewicz et al., 2011). To determine if mutations to the FFD hotspot residues also led to FRQ instability, we analyzed FRQ levels over time by inhibiting translation via the addition of cycloheximide (CHX) and tracking FRQ levels over time (He et al., 2003; Hurley et al., 2013). As seen previously, compared to the half-life of the tagged wild-type strain (FRQ^VHF^), FRQ^FFD2^ showed a decrease in FRQ stability, which may contribute to the low levels of FRQ in the FRQ^FFD2^ strain (Figure 5C and Supplemental Figure 6A) (Guo et al., 2010; Hurley et al., 2013). Conversely and unexpectedly, FRQ^RR/AA^ had a slightly longer half-life than FRQ^VHF^ despite the loss of FRH binding, with FRQ levels that were comparable to the untagged WT strain (Figure 5B and C). In addition to the stability of FRQ in the FRQ^RR/AA^ strain, we also noted a maintenance or increase in the overall levels of the positive transcriptional activator White-Collar 1 (WC-1, NCU02356) (Figure 5D and Supplemental Figure 6B). WC-1 levels are typically lower in strains where *frq* is either not present or not able to interact with the White Collar Complex (composed of WC-1 and WC-2, WCC) to close the feedback loop by mediating the phosphorylation of WC-1 and WC-2 to repress transcriptional activity and stabilize WC-1 (Wang et al., 2019). Together, these data imply that repression via phosphorylation can still occur in the FRQ^RR/AA^ strain without FRH binding, suggesting that FRQ can close the negative feedback loop without binding FRH.

### FRH binding to FRQ is essential for clock robustness, not feedback

As the FRQ^RR/AA^ strain produced stable FRQ that may feedback on the WCC, we tested whether FRQ can close the TTFL independently of FRH. Using a race tube assay to follow overt clock rhythms via the banding phenotype, we found the WT FRQ^VHF^ strain “banded”, whereas the FRQ^FFD2^ strain was arrhythmic, as anticipated (Figure 6A) (Guo et al., 2010). While the FRQ^RR/HH^ strain maintained a typical clock period of ∼21 hrs, the FRQ^RR/AA^ strain had an arrhythmic clock phenotype, correlating with the hypothesis that FRQ/FRH interaction is essential for feedback in the TTFL (Figure 6A).

**Figure 6.**
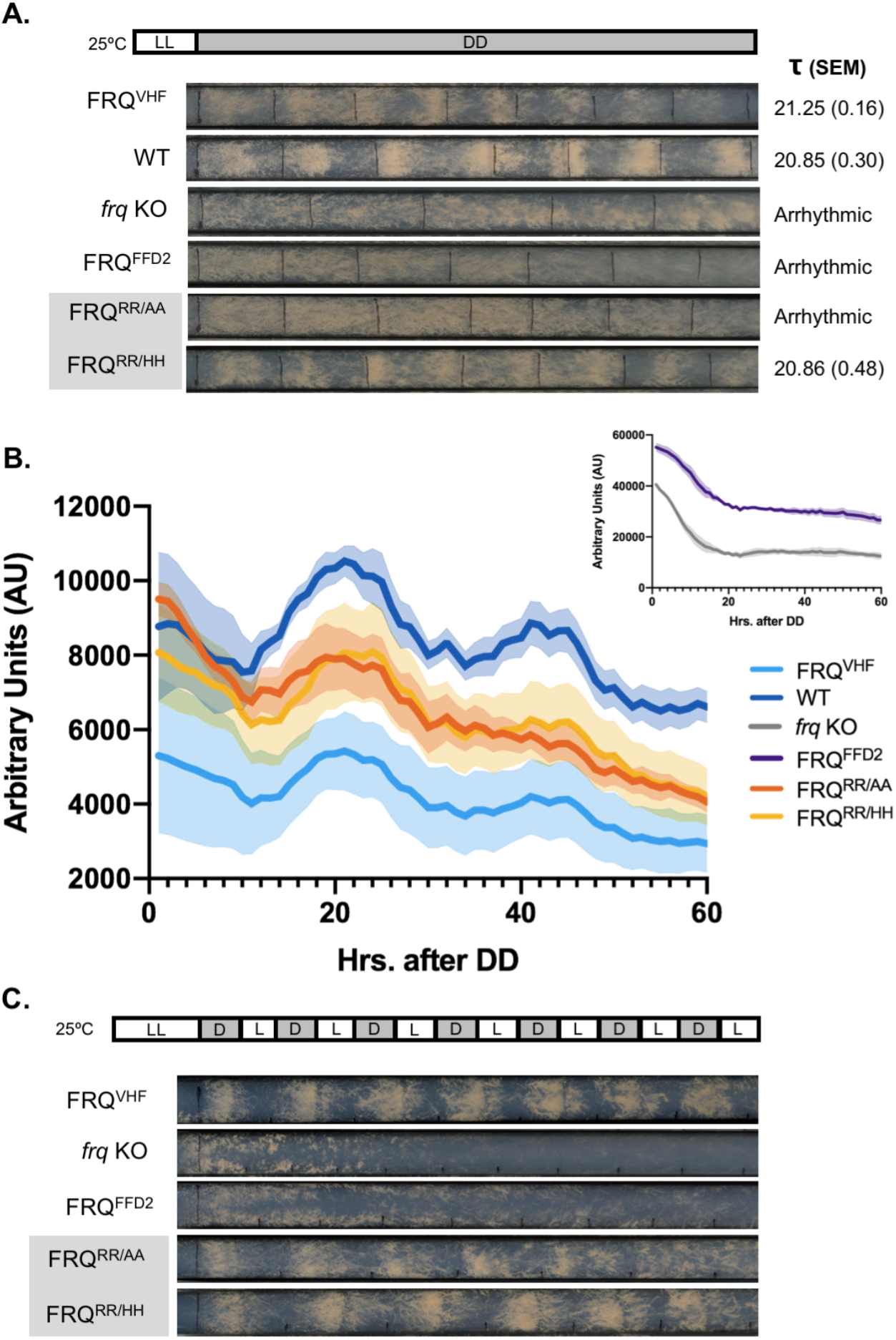
Loss of FRH-FRQ interaction leads to clock dampening rather than arrhythmicity. (A) Representative race tubes of the VHF-tagged FRQ strains described in Figure 5 grown in DD. Average period (**τ**) in hrs. (with SEM) derived from N=5-6 race tubes. (B) Average and standard deviation of luciferase levels over 2.5 circadian days of the VHF-tagged FRQ alleles described in Figure 5 in a strain with a luciferase reporter for *frq* promoter activity (*Pfrq(c-box)::luc*) (N=3) grown in DD in a 96 well format in the presence of luciferin. Inset to separate scales of expression. (C) Representative race tubes of FRQ^VHF^, *frq* KO, FRQ^FFD2^, FRQ^RR/AA^, and ^RR/HH^ strains grown under a 12L:12D lighting regime (N =5-6). Related to Supplemental Figures 6 and 7.

To confirm this clock output phenotype was mirrored in the core clock mechanism, we inserted the same constructs into the *csr-1* locus of an *frq* KO strain harboring a luciferase reporter fused to a minimal *frq* promoter (*c-box::luc*) to assess core clock function, as described above (Bardiya and Shiu, 2007; Gooch et al., 2014). Luminescence output validated that the FRQ^VHF^ and FRQ^RR/HH^ strains maintained circadian oscillations at the level of the TTFL, while the *frq* KO and FRQ^FFD2^ strains showed no oscillations (Figure 6B). Distinctly, the novel FRQ^RR/AA^ strain showed a re-activation of *frq* promoter activity within the first 24 hrs, which dampened on the second day, consistent with FRQ^RR/AA^ closing the feedback loop but not robustly reactivating the TTFL.

To corroborate FRQ^RR/AA^ was able to close the feedback loop, we assessed conidiation in race tubes in a 12hr light:12hr dark (12L:12D) entrainment cycle, which will only oscillate if FRQ is able to close the feedback loop. While FRQ^VHF^ and FRQ^RR/HH^ strains showed banding rhythms, and the FRQ^FFD2^ and *frq* KO strains were arrhythmic (Figure 6C), the FRQ^RR/AA^ strain showed banding rhythms (Figure 6C). Taken together, this data is consistent with the ability for FRQ^RR/AA^ to repress the WCC without FRH, which is directly in contrast to the prevailing theory that FRH is necessary for FRQ to close the negative feedback loop (Cheng et al., 2005; Shi et al., 2010).

## Discussion

An understanding of how protein-protein interactions underly the mechanisms that regulate circadian feedback and post-transcriptional regulation has been hampered by the highly disordered nature of the core clock proteins. To investigate these protein-protein interactions, we adapted an approach we termed LOCATE, which employs immobilized synthetic peptide microarrays to capture interactions across the full length of proteins enriched in IDRs (Clerc et al., 2021; Ivarsson and Jemth, 2019; Katz et al., 2011). Applying LOCATE to the *N. crassa* disordered negative-arm protein FRQ, we identified a SLiM hotspot beyond the genetically-identified binding region for FRH on FRQ, while also uncovering novel sites of FRH interaction. These multivalent interactions support a model of a fuzzy negative arm protein complex mediated by both a specific SLiM and non-specific electrostatic interactions (Figure 7A).

**Figure 7.**
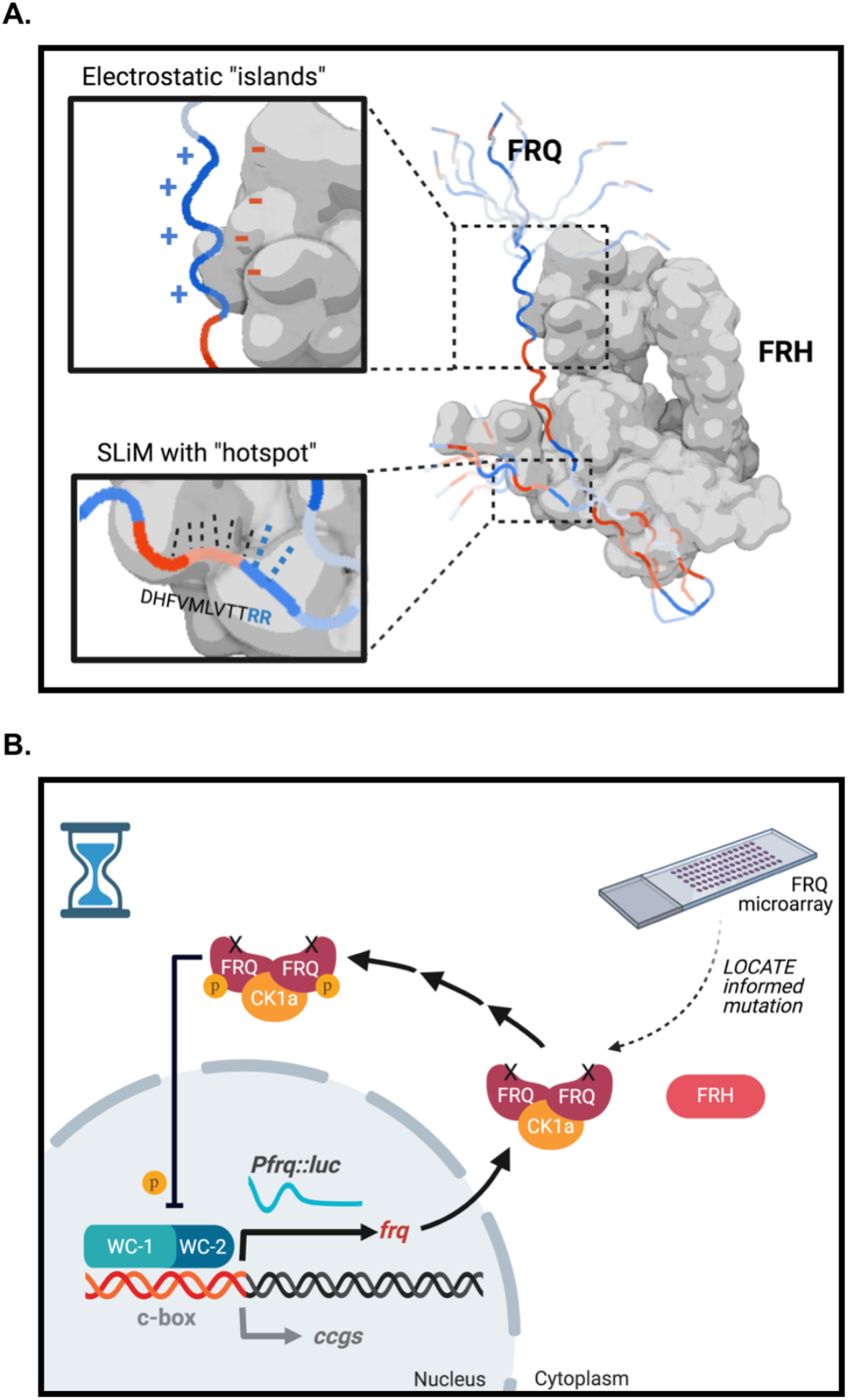
Proteins in the negative arm of the clock form a multivalent complex aided by electrostatic islands and a SLiM hotspot within FRQ that is important for clock robustness. (A) A space-filling model of the known crystal structure of FRH (PDB 4XGT; grey protein) and a model of a portion of FRQ (linear protein) using AlphaFold2 (Jumper et al., 2021), not shown to scale. Blue regions represent positively charged islands and red regions represent negatively charged islands. The top inset shows expected electrostatic interactions between positively charged islands on FRQ and the overall negative outer surface of FRH. The bottom inset shows the specific FFD motif, with the double arginine hotspot highlighted in blue. FRQ = FREQUENCY, FRH = FRQ-Interacting RNA Helicase. (B) Model based on the LOCATE-informed mutation. FRQ^RR/AA^ is not able to interact with FRH yet can feed back on WC-1/2 to close the circuit in an hourglass-like manner. The loss of robustness in the clock is demonstrated by the damping oscillation of luciferase as expressed from the *frq* promoter. Abbreviations as in Figure 1.

The high amino acid resolution of the LOCATE approach allowed for pinpoint recognition of a hotspot, two novel arginine residues near the FRQ-FRH binding domain (FFD), that is critical for FRQ/FRH interaction *in vitro* and *in vivo*. This RR783AA region was unique in that, while breaking the binding between FRH and FRQ when mutated, it did not cause FRQ to become unstable (Figure 5B and C). This was unexpected, as it has been noted that previous mutants that affect FRQ/FRH interaction also affect FRQ stability (Cheng et al., 2005; Guo et al., 2010; Hurley et al., 2013). This instability has led to the hypothesis that the role of FRH is to act as a Nanny for FRQ, stabilizing FRQ so it can complete the feedback loop (Hurley et al., 2013). How FRH interaction enhances stability is not known, but was hypothesized to occur through the shielding of regions of FRQ that enhance rates of degradation (Hurley et al., 2013). Our data suggests instead that the FFD region may encode two distinct modes; one that binds FRH, and a second that is recognized for protein degradation (i.e. a degron) (Ravid and Hochstrasser, 2008). In this model, mutations that disrupt the double arginine impede both FRH and degron recognition, whereas FFD variants that retain the arginine residues disrupt FRH binding but are still recognized and degraded. This degron hypothesis will be a target of future investigation.

Given that LOCATE allowed us to identify specific residues which, when mutated, disrupted FRQ/FRH interaction without leading to FRQ degradation, we were able to investigate the role of FRQ in the core clock independent of the FRQ/FRH interaction. While the clock was not able to robustly run in the absence of FRQ/FRH interaction, FRQ was able to exert negative feedback on to the WCC, letting the clock run in an hourglass-like manner (Figure 7B). In concordance with this data, FRQ can interact with the WCC and its principle kinase Casein Kinase 1a (CK-1A) independently of FRH, enter the nucleus without FRH, and decreased binding of FRH to FRQ can maintain clock rhythmicity (Cha et al., 2011; Conrad et al., 2016; Froehlich et al., 2003; Guo et al., 2010; Hurley et al., 2013; Shi et al., 2010). This evidence supports a model where FRQ and CK-1A can carry out WCC repression at the *frq* promoter without the input of FRH, suggesting that contrary to the current paradigm, FRH is essential for robustness but not for feedback (Figure 7B).

How FRH may impart such robustness is unclear but could occur through multiple, non-mutually exclusive, avenues. FRH localizes FRQ to the cytosol after WCC repression, which could tune WCC activity (Cha et al., 2011). FRH may also regulate the phosphorylation rate of FRQ, a parameter that is important for self-sustained oscillations, though the ATPase function of FRH is not necessary for clock function (Hurley et al., 2013; Lauinger et al., 2014; Upadhyay et al., 2020). In higher eukaryotes, another negative arm protein, CRYPTOCHROME (CRY), contributes to clock robustness rather than feedback (Putker et al., 2021). These data highlight the need to investigate which proteins are essential for the TTFL and the role of robustness in circadian timing.

Beyond the specific FRQ/FRH FFD region/hot spot, LOCATE highlighted regions of FRQ whose interaction with FRH was principally electrostatically driven by positively charged islands (Figures 2 and 3A). The presence of these positively charged islands was conserved across FRQ orthologs in fungi and higher eukaryotes (Figure 3C and Supplemental Figure 3A) and maintained upon the phosphorylation of over 80 sites that occur on FRQ (Supplemental Figure 2C) (Baker et al., 2009). Modulation of one of these islands affected the function of the clock in *N. crassa*, leading to a severe increase in oscillatory period (Figure 3E). It is noteworthy that previously identified point mutations in *frq*, such as *frq*^3^ (E364K) and *frq*^7^ (G459D), both involving substitutions to charged residues, have yielded increases in clock period (Aronson et al., 1994).

The presence of conserved positively charged islands on FRQ, and their role in FRH-binding, parallels work illustrating the importance of electrostatic interactions and multivalency in mammalian clock proteins and in other complexes comprising disordered proteins (Czarna et al., 2011; Holehouse, 2019; Ozber et al., 2010; Sherry et al., 2017; Tompa and Fuxreiter, 2008; Xu et al., 2015; Zhao et al., 2021). This pattern of multivalent electrostatic islands supports a fuzzy interaction model, where the proteins interact as a heterogeneous ensemble rather than a single fixed structure (Buljan et al., 2012; Sharma et al., 2015; Tompa and Fuxreiter, 2008; Wiggers et al., 2021). Relevantly, electrostatic interactions can increase the binding affinity between proteins in a fuzzy complex (Bugge et al., 2020; Shen et al., 2018; Wiggers et al., 2021). This fuzzy complex model is further supported by evidence of a change in the conformational dynamics of the negative arm protein complex over the circadian day (Pelham et al., 2021). We therefore propose that the FRQ/FRH complex meets the definition of a fuzzy complex, although further investigation of the conformational dynamics of this interaction is warranted (Tompa and Fuxreiter, 2008). This combination of positively charged residues that can work in concert with hydrophobic residues is reminiscent of acidic activation domains, where negatively-charged residues conspire with hydrophobic sidechains to determine specificity and affinity (Erijman et al., 2020; Ravarani et al., 2018; Sanborn et al., 2021; Staller et al., 2021, 2018; Tuttle et al., 2018). Whether or not these modes of interactions differ only in terms of the charge sign, or if additional chemical and structural nuances emerge, remains an area of open investigation.

As the electrostatic context and disorder surrounding a motif affects SLiM binding, and many FRQ SLiMs occurred near negatively charged islands, it is tempting to speculate that FRH electrostatically steers the availability of non-FRH SLiMs on FRQ (Figure 7A and Supplemental Figure 2D) (Bugge et al., 2020; Das and Pappu, 2013; Ganguly et al., 2012; Ivarsson and Jemth, 2019; Tompa and Fuxreiter, 2008). Electrostatic steering could regulate protein-protein interactions to time the formation of unique, or temporally-specific, negative arm macromolecular complexes (Pelham et al., 2021). Considering the conservation of positively charged islands as a molecular feature in negative arm IDRs in higher eukaryotes, it is possible this is a widely conserved mode of regulation in many clock proteins (Pelham et al., 2020; Zarin et al., 2019). Given the insights garnered from the investigation of FRQ/FRH interaction, we propose that the LOCATE method could also be employed broadly to investigate interactions between highly disordered clock proteins, perhaps focusing on the effect of post-translational modifications (Dunker et al., 2005; Baker et al., 2009; Mosier and Hurley, 2021; Blikstad and Ivarsson, 2015; Dittmar et al., 2019; Tapia et al., 2008; Pelham et al., 2021, 2020). As clock protein IDRs, and IDRs in general, are potentially druggable targets, the LOCATE method may be relevant to identify candidate SLiMs within IDRs for drug development (Katz et al., 2011; Van Roey et al., 2014; Wójcik et al., 2018).

## Acknowledgements

We thank the Fungal Genetic Stock Center and the Dunlap and Loros labs at Dartmouth College for providing *Neurospora crassa* strains. Lab technical support was kindly provided by Christopher Kirchhoff, Ellinor Tai, and Samantha Keller. We thank the RPI Core Facilities for the use of equipment, particularly Dr. Sergey Pryshchep and the Cell and Molecular Biology Research Core. Figures 1A, C, D, and 7, and Supplemental Figure 2D, were created using Biorender.

## Author Contributions

Conceptualization, J.M.H.; Methodology, M.S.J., D.G., D.G.S., J.F.P., P.K., A.S.H., J.M.H.; Investigation, M.S.J., D.G.S., J.F.P., J.T.; Formal Analysis, M.S.J., D.G., D.G.S, J.F.P, G.M.G., J.T.; Software, D.G., G.M.G., A.S.H.; Resources, P.K., A.S.H., J.M.H.; Writing – Original Draft, M.S.J., D.G., J.F.P.; Writing – Review & Editing, M.S.J., D.G., D.G.S., J.F.P., A.S.H., J.M.H.; Visualization, M.S.J., D.G., D.G.S., J.F.P.; Supervision, M.S.J., P.K., A.S.H., J.M.H.; Funding Acquisition, M.S.J., D.G., D.G.S., A.S.H., J.M.H.

## Methods

### Peptide library design, synthesis, and microarray printing

Two peptide libraries were designed, with Library I primarily made up of a linear peptide “mapping” of the primary sequence of FRQ (NCU02265) and Library II containing further rationally designed peptides to investigate the specificity of FRH binding to original “parent” peptides (Supplemental File 1). The FRQ “mapping” peptides in Library I were 15 amino acids (a.a.) in length, beginning at the N-terminus of FRQ’s sequence, with consecutive peptides shifting by 3 a.a.. Peptide mapping for some regions of interest (eg. FFD region) were repeated at a finer scale of 15 a.a., shifting by 1 a.a. between consecutive peptides. Any parent peptides of interest were used as the basis of further rationally designed peptides, such as scrambled sequences (using the Genscript random library tool; https://www.genscript.com/random_library.html) or truncation series, to identify sequence-specific interactions and the minimal SLiM. Parent peptides containing a candidate SLiM were further singly or doubly mutated to other residues of interest (not all combinatorial possibilities were investigated) to identify permissive or prohibited residues at different positions within the SLiM.

Peptides were synthesized using standard Fluoroenylmethyloxycarbonyl (Fmoc) chemistry in an automated peptide synthesizer (Multipep RS, INTAVIS Bioanalytical Instruments AG, Germany), as done previously (Shastry and Karande, 2019). Specifically, parallel peptide synthesis proceeded from C- to N-terminus on solid cellulose discs, with all peptides N-terminally acetylated to better mimic the charge of a peptide segment within the parent protein’s sequence. Note that the first peptide based on the parent protein’s N-terminus was not N-terminally acetylated. See Supplemental File 1 for more details on the two peptide libraries. Once synthesized, peptide-cellulose discs were reconstituted in 250 µL dimethyl sulfoxide (DMSO) following Intavis’ standard work-up procedure. The resulting peptide stock solution were used 1:1 for spotting in triplicate in a microarray format on nitrocellulose-coated glass microscope slides using a slide-spotting robot (Intavis Bioanalytical Instruments AG). Peptide microarrays were then air-dried for 2 hrs at 65°C.

### FRH Protein Expression and Purification

The FRHΔ100 plasmid (Conrad et al., 2016), consisting of a pET28a vector, FRH (100-1106 a.a.), and an N-terminal 6x His tag, was transformed into BL21 (DE3) Competent *E. coli* cells (New England Biolabs, C2527) and plated on selective LB plates containing 30 µg/ml kanamycin (AMRESCO, 0408-10G). Positive colonies were grown in 3L of liquid LB with 30 µg/ml kanamycin at 37°C, 225 rpm, until Abs_600_ was ∼0.45. After cooling the culture on ice, protein expression of the FRHΔ100 plasmid was induced with 0.2 mM IPTG (Biotium, 10021), at 18°C, 185 rpm, for 16 hrs. Cells were pelleted by centrifugation at 4°C, 4800 rpm, for 15 minutes, and kept on ice for immediate protein extraction. Protein extraction buffers contained 50mM HEPES (Sigma Aldrich, H0887-100 ml) and 150 mM NaCl, pH 7.0, and varied amounts of imidazole (Amresco, 0527-100G). Pelleted cells were resuspended in 30 ml of Lysis Buffer that contained 10 mM imidazole. Cells were lysed by three rounds of French Pressing at 1000 psi. The cell lysate was clarified by centrifugation at 4°C, 14,000 rpm for 20 min. The soluble fraction was split between two columns each with 2 ml Ni-NTA agarose beads (Qiagen, 30210) pre-equilibrated with Lysis Buffer, and nutated for 1 hr. at 4°C. The columns were washed with two rounds of 15 ml Wash Buffer containing 30 mM Imidazole. Proteins were eluted in 500 µL fractions using an Elution Buffer with 200 mM imidazole. Each elution was applied to a 40 kDa cut-off Zeba column (ThermoScientific, 87769) to de-salt and exchange the buffer to PBS (100 mM NaCl, 10 mM Potassium Phosphate, pH 7.0). A BSA-based Bradford assay (BioRad, 5000006) was used to quantify resulting amounts of FRH protein, and the percentage of FRH in the final product was visually estimated based on the percentage of the FRH band relative to other bands (∼30% FRH) when the eluted protein was visualized on a Coomassie-stained gel (Amresco, 0472-25G). Expression and purification of FRHΔ100 was verified by SDS-PAGE analysis (ThermoScientific, WG1602BOX), and Western blotting with primary anti-His at 1:1000 (Sigma Aldrich, SAB1305538), secondary anti-Mouse at 1:10000 (ThermoFisher, 31430) and Pico (ThermoScientific, 34577). Further purification was not considered as the non-specific protein products acted beneficially as a built-in competition assay to decrease false positives for FRH binding.

### Microarray incubation and data analysis

Basic microarray screening protocol was carried out at room temperature on a rocker using 5 mL of each buffer per slide, as in (Shastry and Karande, 2019). FRQ-based microarrays were first blocked for 3 hrs in PBS (10 mM phosphate, 100 mM NaCl, pH 7.0) with 5% w/v BSA, followed by 3 x 10min washes in PBS. Incubation with different approximate concentrations (1, 10, or 100 nM) of the purified and buffer-exchanged FRH occurred for 3 hrs followed by another 3×10 min washes and 1 hr. incubation with antibodies (in 2.5% w/v BSA in PBS) with 3×10 min washes between each step, either anti-His at 1:1,000 then anti-Mouse at 1:500,000 (see above), or anti-FRH at 1:12,500 (Shi et al., 2010) followed by anti-Rabbit at 1:500,000 (Invitrogen, 31460), then SuperSignal West FEMTO (ThermoScientific, 34094). Non-specific antibody binding was assessed by incubating with antibodies and FEMTO, but without first incubating with FRH. Chemiluminescence imaging was performed with a ChemiDoc XRS+ System (Bio-Rad) and Image Lab 4.0 software, using signal accumulation mode (SAM) with the high-resolution option with 2×2 binning.

Microarray images were normalized for each library within ImageLab by standardizing the intensity range to allow better comparison amongst replicates, and exported to Fiji (ImageJ v2.0.0, NIH) to convert to 8-bit grayscale, inverted, and background subtracted (rolling ball radius = 50 pixels) (Rueden et al., 2017). TIGR Spotfinder (Release 2009-08-21) was used to quantify spot intensities using the Otsu segmentation method with local background subtraction (Saeed et al., 2003) (Supplemental File 1). Basic plots of sequential normalized microarray intensity, truncations, and scrambled peptides were all plotted in PRISM 9.0.2. Compositional analysis of the peptide residues was also carried out as in (Shastry and Karande, 2019), and plotted in PRISM 9.0.2.. The frequency of occurrence of residue types (acidic, basic, polar, aromatic, or nonpolar) in the 3-mer shift peptide mapping portion of the peptide library were compared to the frequency of occurrence in the top 10% of peptides, i.e., the peptides with the highest normalized binding intensities. The statistical significance of the changes in residue occurrence amongst populations was determined using the test statistic z:

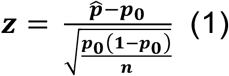

where ***p̂*** is the percentage of a residue or residue type occurring in the top 10% of peptides; ***p*_0_** is the percentage of the same residue or residue type occurring in the peptide mapping portion of the library; and ***n*** is the total number of residues in the partial peptide mapping library. For net charge at pH 7.0 calculations, residues D and E were each considered −1 charge, R and K as +1 charge and H as +0.091, while all other residues were considered charge neutral. Resulting peptide charges were plotted in PRISM 9.0.2. Analysis for the FFD region was calculated as a fraction of the WT peptide LPDDHFVMLVTTRRV value of 64, an average of the Library II peptides 215 and 235 (Supplemental File 1), that was then transformed to the log_2_ scale.

### Electrostatic potential calculation for FRH

The solved crystal structure of FRH was downloaded from the Protein Data Bank (PDB 5E02; Conrad et al., 2016). Using PyMol (v2.4.0), the file was converted to PQR format and the APBS tool used to calculate the solvent-accessible electrostatic potential according to the Poisson-Boltzmann equation (Fogolari et al., 2002). The electrostatic map was visualized in PyMol, using a gradient of −3 kT/e (red) to +3 kT/e (blue).

### Bioinformatics for BLAST Multiple Sequence Alignment and Cladogram

The *N. crassa* FREQUENCY (NCU02265) protein sequence was run in NCBI BLAST (Sayers et al., 2021). Homologous protein sequences with over 50% identities or 50% positives with FRQ were chosen for further analysis, yielding a list of ten other homologous proteins. Fungal protein sequences of interest were exported from FungiDB, release 47 (Basenko et al., 2018). Next the protein sequences were aligned using the UniProt alignment tool and exported in Stockholm format (The UniProt Consortium, 2019). The multi-sequence alignment was further visualized in SnapGene (version 5.1.1) and a logo was made using the interactive tool Skylign (accessed July 2020). Parameters used in Skylign were logos based on the full alignments, letter heights determined by information content above background, and their weighted counts method, where weights are applied to account for highly similar sequences before calculating a maximum-likelihood estimate for each column in a multi-sequence alignment (Wheeler et al., 2014). The relationship amongst the species included in our FRQ homology analysis were represented in a rectangular cladogram, created using the Interactive Tree of Life online tool, ver. 4 (Letunic and Bork, 2019).

### Generation of FRQ^VHF^ and FRQ-FFD mutant strains

An FRQ^VHF^ cassette was developed for insertion into the cyclophilin locus of *N. crassa* as described (Bardiya and Shiu, 2007). Starting at the 5’ end, we fused 1000 bp from the upstream portion of the cyclophilin locus (*csr-1,* NCU00726) from FungiDB to the 3000 bp upstream promoter of the *frq* ORF and the complete wildtype *frq* Open-Reading Frame (ORF), followed by a 10x glycine linker and V5-10His-3Flag (VHF) tag (Emerson et al., 2015), followed by a new stop codon, followed by 1000 bp downstream of the *frq* ORF, and finally 1000 bp from the downstream portion of the target csr-1 locus (Basenko et al., 2018). Template genomic DNA was harvested from a wildtype *N. crassa* strain 87-3 (bd+, mat a) using the Gentra Puregene Tissue kit (Qiagen, 158622) and following the manufacturer’s instructions. PCR reactions using primers found in Table 1 were carried out using Phusion Flash High-Fidelity PCR Master Mix (ThermoFisher, F548S), following the manufacturer’s instructions and using an Eppendorf Mastercycler Nexus Thermal Cycler with the following program: 98°C 10 sec, (98°C 5 sec, 67°C 5 sec, 72°C 4 min) x35, 72°C 5 min, 4°C 5 min. Appropriate cDNA product sizes were confirmed using a 0.8% agarose gel in 1X TAE buffer, relative to a 1 kb DNA ladder (New England Bio, N3232L). The cDNA pieces with primer overhangs were ligated along with a gapped selective yeast vector (pRS426, containing *URA3* and ampicillin resistance; Hurley et al., 2013) using the homologous recombination system endogenously found in yeast (strain FY834 from the Fungal Genetics Stock Center), by carrying out a Lithium Acetate/ PEG transformation (Collopy et al., 2010; Gietz and Schiestl, 2007). After isolating all plasmid DNA using a “Smash and Grab” protocol (Hoffman, 2001), the harvested plasmids were transformed into *E. coli* (DH5alpha derivative; New England Bio, C2989K) using the manufacturer’s instructions (BTX Harvard Apparatus, 45-2001) and plated onto LB agar plates with 100 µg/ml ampicillin to grow overnight at 37°C. Colonies were picked and grown in liquid LB broth with 100 µg/ml ampicillin overnight at 37°C and shaken at 225 rpm. After centrifugation of the culture, plasmids were isolated and purified using the QIAprep Spin Miniprep kit (Qiagen, 27106), and PCR of the final linear cassette was done as stated above except using primers MSJ001F and MSJ006R. The cassette was purified using the QIAquick PCR Purification Kit (Qiagen, 28106). Cassette sequences were verified through Sanger sequencing using sequencing primers spaced ∼700-800 nt apart (Genewiz and Eurofins Operon, LLC). Verified cassettes were transformed into the *csr-1* locus of an *frq* KO *N. crassa* strain, 122 (bd+, delta-frq::hph+, mat a), using methods previously described (Bardiya and Shiu, 2007). Transformants recovered after electroporation in a liquid VM medium with additional yeast extract and were then plated on an agar plate containing a growth-restrictive mixture of fructose-glucose-sorbose (FGS) and 5 µg/ml of cyclosporin A (Bardiya and Shiu, 2007; Park et al., 2011). After 3-4 days of growth at 25°C, isolated colonies were picked and placed onto selective slants (small test tubes with agar-based VM media) and 5 µg/ml cyclosporin A (Sigma, 20024).

We designed our FRQ mutations around the concept of maintaining the relative adaptiveness of codons used, since previous studies have shown that FRQ codon usage has effects on its structure and stability (Zhou et al., 2013). By accessing the codon table for *N. crassa* (Nakamura et al., 2000), we calculated the relative adaptiveness of each codon, with 100% being the highest frequency amino acid codon, and then relatively scaled the frequency of the remaining codons (Fuhrmann et al., 2004). We used this Relative Adaptiveness measure of codons to rank from most adaptive to least adaptive and substituted an alanine or histidine codon of similar rank for our mutations. For our FRQ-FFD mutants, the program PrimerX (http://www.bioinformatics.org/primerx/cgi-bin/DNA_1.cgi) was used to design the needed primers, using the following specifications: Melting temp. 50-85°C, GC content 40-60%, length 40-60 bp, 5’ flanking region 20-30 bp, 3’ flanking region 20-30 bp, terminates in G or C, mutation site at center, and complementary primer pair (Lapid and Gao, 2006). Designed primers are found in Table 1.

**Table 1.**
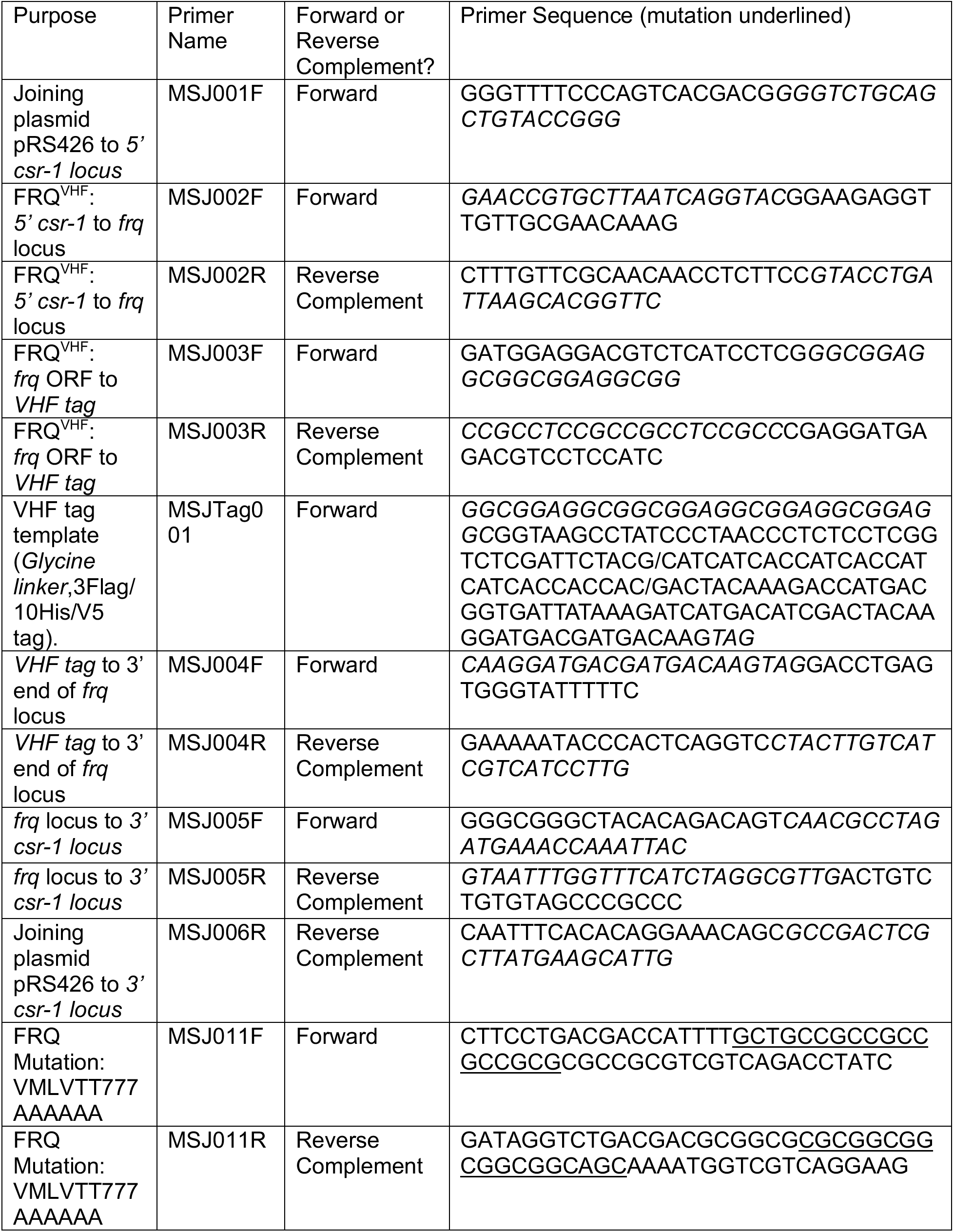
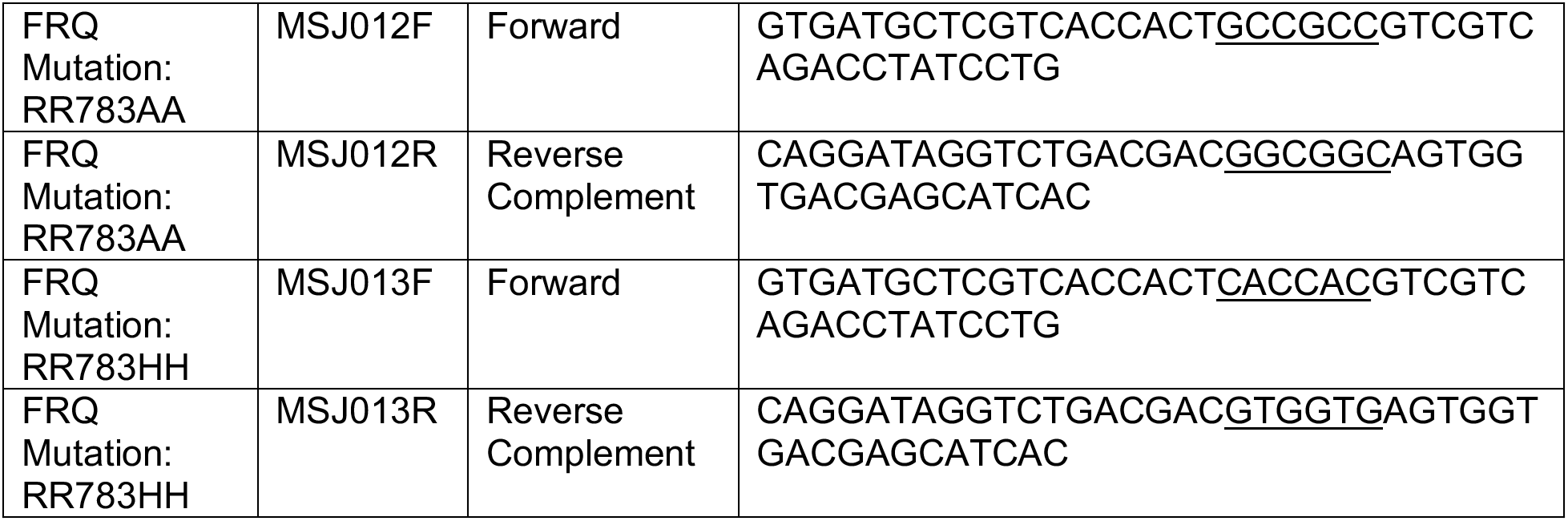
Primers used in this study synthesized from Integrated DNA Technologies, Inc. IDT. Underlined bases denote the substitutions made for the different FRQ mutations.

For luciferase reporter assays, we created a new strain (uber #6) which was the result of a cross between the X200-3 strain (bd+, his3::pLL26, mat A) where a minimal frq promoter fused to luciferase (Pfrqmin1::luc), and the 122 strain (*frq* KO), to create an *frq* KO strain with a luciferase reporter attached to a minimal frq promoter in the background (Delta-frq::hph+, bd+, his3+::Pfrqmin::luc). For the race tube and Co-IP assays, we used WT strain 328-4 (bd+, mat A).

### Race tube assays and period determination

Race tube assays were carried out using custom glass tubes filled with 14 mL of race tube medium (1x Vogel’s salts, 0.05% glucose, 0.1% arginine, 50 ng/ml biotin and 1.5% bacto-agar). See fgsc.net/neurosporaportocols/How to choose and prepare media.pdf for more details on Vogel’s salts or other Neurospora media. Prepared race tubes were inoculated with 10 µL of a conidial suspension from the noted strain and grown for ∼24 hrs at 25°C in constant light, before being synchronized with a transition to constant dark, 25°C, and marked each 24hrs until strains reached the end of the race tube. Race tubes were scanned using an EPSON GT-1500 scanner, and images were cropped and converted to black and white. Clock period was analyzed using ChronOSX (v1.1.0) (Roenneberg and Taylor, 2000), using the Period Analysis option and including 6 days of constant dark densitometry data from each race tube to calculate the mean and standard deviation for each tube, followed by a mean and standard error of the mean for each set of replicates per strain.

### CCD array trials and analysis

Low Nitrogen-CCD media (LN-CCD; 0.03% glucose, 0.05% arginine, 50 ng/ml biotin, 1x Vogel’s salts, 1.5% bacto-agar, 25 µM luciferin) with 0.001 M Quinic Acid (pH 4.75) was used for all luciferase reporter assays. 185 µL of this media was used per well in a black 96-well plate (Eppendorf, 951040196), inoculated with 10 µL of the relevant conidial suspension. Plates were sealed with a breathable membrane (BreatheEasy, USA Scientific, 9123-6100), and incubated at 25C in constant Light for ∼48 hrs before being placed at 25C constant darkness in an incubator with a PIXIS CCD array (Princeton Instruments, 1024B) with a 35 mm Nikon DX lens (AF-S NIKKOR, 1:1 8G), run by the program Lightfield (version 5.2, Princeton Instruments). Images were acquired for 15 min every hour and final image stacks were imported into FIJI (ImageJ v2.0.0, NIH) to adjust brightness, and denoised using the default “Remove outliers” tool option. A custom image analysis plugin was used called “Toolset Image Analysis Larrondo’s Lab 1.0” (courtesy of Luis Larrondo) using the 96-well plate quantifying tool. Data were smoothed using a moving average of three timepoints, before plotting in PRISM 9.0.2.

### Co-immunoprecipitation and western blotting

Tissue from each *N. crassa* strain was grown by making a conidial suspension using 1mL of Liquid Culture Media (LCM; 2% Glucose, 0.5% Arginine, 1x Vogel’s Salts, 50ng/ml Biotin) to resuspend conidia from a ∼1 week old Vogel’s minimal media slant. The centrifuged and washed conidia were then inoculated into a 125 mL flask containing 50 mL of LCM and grown at 25°C, 125 rpm in constant light for 48 hrs before harvesting. Harvesting was carried out using vacuum filtration and flash freezing in Liquid Nitrogen before storing at −80°C. Tissue was ground in a mortar and pestle along with Liquid Nitrogen, before extracting tissue lysate using a similar volume of chilled Protein Extraction Buffer was added (pH 7.4, 50 mM HEPES, 137 mM NaCl, 10% Glycerol, 0.4% NP-40 alternative) with 1x HALT protease and phosphatase inhibitor, EDTA-free (87785, ThermoScientific). Co-Immunoprecipitation employed 40 µL of resuspended anti-FLAG magnetic beads (M8823, Sigma) or anti-V5 agarose beads (A7345, Sigma) that were prepared according to manufacturer’s instructions for each 3.5 - 4.8 mg of total lysate, as measured by Bradford assay. Final volumes were brought up to 1 mL total with further Protein Extraction Buffer with 1x HALT. Samples were incubated overnight at 4°C while nutating. Following placement on a magnetic rack, the flow through was removed and the beads were washed three times with 1 µL of Protein Extraction Buffer. Eluted proteins were retrieved by adding 40 µL of 2x LDS Buffer (Cat #) and boiling for 15 min. at 65°C. Samples were removed to a fresh Eppendorf tube and boiled at 100°C for 5 min. with the addition of 3%v/v ß-mercaptoethanol before freezing at −20°C. Thawed samples were run on precast NuPAGE 3-8% Tris-Acetate gels (Invitrogen, WG1602BOX) following manufacturer’s protocol, and transferred to PVDF membrane using a BioRad Trans-Blot Turbo Transfer System (BioRad, 1704150). Membrane was blocked using 5% Milk in PBS buffer with 0.2% Tween-20, Primary antibodies were custom antibodies, courtesy of the Dunlap-Loros Labs at Dartmouth, used at 1:5000 for anti-FRQ (Garceau et al., 1997), 1:12000 for anti-FRH (Shi et al., 2010) or 1:5000 for anti-WC1 (Lee et al., 2000), in 1% Milk with PBS and 0.2% Tween-20. Secondary antibody was 1:5000 (Goat anti-Rabbit; Invitrogen, 31460) in PBS and 0.2% Tween-20. Western blots were incubated with SuperSignal West FEMTO (ThermoScientific, 34094) or ATTO (ThermoScientific, A38554) for anti-WC-1 blots, and imaged using a Bio-Rad GelDoc imager.

### Cycloheximide assay and semi-quantitative western blotting

The cycloheximide assay was adapted from Hurley et al. (Hurley et al., 2013). Briefly, conidial suspensions from the designated *N. crassa* strain were inoculated into a petri dish filled with LCM (2% glucose) and grown in constant light at 25°C. After 24-36 hours (dependent upon the growth rate of the strain) plugs were cut from the resulting mycelial mat and placed into individual Erlenmeyer flasks with ∼50 ml of LCM (2% glucose). After plugs grew in constant light at 25°C (125 rpm) for a further 24 hours, 40 µg/uL of Cycloheximide (94271, VWR) was added to each culture and this was designated time 0. Samples continued to grow at LL, 25°C, 125 rpm until they were harvested using vacuum filtration at different timepoints (0, 2, 3, 4, 5, 6, 7 hrs post-cycloheximide addition) via flash freezing in liquid nitrogen before storing at −80°C. Protein extraction and standardization was executed as described above. SDS-PAGE and Western blot protocols were carried out as above, except using precast NuPAGE 4-12% Bis-Tris gels (Invitrogen, WG1402BOX), 1:5000 anti-V5 primary (Invitrogen, 46-1157) and 1:25000 Goat anti-Mouse (Invitrogen, 313430) secondary antibodies, and SuperSignal West FEMTO (ThermoScientific, 34094) or ATTO (ThermoScientific, A38554) in the case of FRQ^FFD2^. Image Lab software (version 6.0.1) was used for relative quantification and Amido Black staining was used as a total protein loading control for normalization. Pixel density was quantified from hour 0 to 7 or 0 to 3 hours within each western blot (dependent upon the stability of FRQ) and normalized by dividing by the pixel density of a matching area of the Amido Black stained membrane. Normalized data was plotted as a ratio relative to timepoint 0 in PRISM 9.0.2.. An exponential fit was made to the biological triplicate timepoints for each strain, and a half-life calculated using first-order decay kinetics (Hong et al., 2008; Hurley et al., 2013). For the WC-1 relative lysate levels, pixel density after normalization by the matching area of the Amido Black stained membrane, was plotted as a ratio relative to FRQ^VHF^ WC-1 levels, again plotted in PRISM 9.0.2..

### Sequence analysis

Overall predictions of FRQ sequence disorder were carried out using the package metapredict, a machine learning-based method that predicts whether a residue is in a disordered region by predicting that residue’s consensus score (smoothing using a 5 residue window) across multiple disorder predictors (Emenecker et al., 2021). Metapredict considers residues with a predicted consensus value of >0.3 as disordered (meaning that more than 30% of the predictors agreed that the residue was in a disordered region), while residues <0.3 are considered ordered. To calculate sequence conservation, we extracted 86 orthologous *N. crassa* FRQ sequences from the eggNOG (v5.0) (Huerta-Cepas et al., 2019), aligned these sequences, then calculated per residue conservation scores by using the approach described by (Capra and Singh, 2007), grouping amino acids by residue properties. Using the primary amino acid sequence of FRQ (NCU02265), we calculated the net charge per residue (NCPR) using a sliding window of 15 residues and average Kyte-Doolittle hydropathy using the localCIDER program (v0.1.18) with a sliding window size of 5, meaning the value at each position is based on that residue and the two residues to each side (Holehouse et al., 2017). NCPR was also recalculated at different timepoints including the phosphorylations reported in Baker et al. (2009) as negative charges, and plotted using PRISM 9.0.2. We measured clustering of particular residue classes (*e.g.*, positive, negative, and aromatic residues) within FRQ’s sequence by calculating the average inverse weighted distance (IWD). The IWD is defined as:

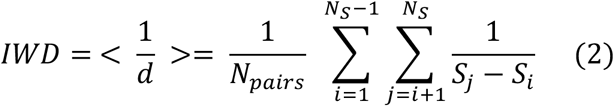

where *S* is the set of target residue positions, *S_i_* is the i-th element of *S*, *N_S_* is the number of items in *S*, and *N_pairs_* is the number of pairwise combinations between elements of *S* (Schueler-Furman and Baker, 2003; Yang et al., 2020). The IWD was then compared to the IWD calculated from 10,000 randomly shuffled FRQ sequences to assess significance as IWD values above the 95 percentile, and following the approach presented in (Cohan et al., 2021) we calculated a residue class specific Z-score to allow comparisons across orthologs that differ in length and amino acid composition.

### All-atom simulations

All-atom simulations were performed using the CAMPARI simulation engine with the ABSINTH implicit solvent model (Vitalis and Pappu, 2009a, 2009b) (http://campari.sourceforge.net/). A 50-residue fragment from FRQ (residues 754 – 803) was simulated in a spherical droplet with a radius of 94 Å. Simulations were performed at 340 K and 15 mM NaCl as has been done previously (Cubuk et al., 2021; Martin et al., 2020; Sherry et al., 2017). Ten independent simulations were run for each of the different constructs, which include a wildtype construct, an RR to AA construct, an RR to HH construct with neutral histidine, and an RR to HH construct with a positively charged histidine. Each simulation was run for 126 x 10^6^ Monte Carlo steps, with the first 6 x 10^6^ discarded as equilibration. Trajectory information was saved every 80,000 steps, such that the final ensembles consist of 15,000 distinct confirmations.

Simulations were analyzed using MDTraj and SOURSOP (https://soursop.readthedocs.io/). Secondary structure was calculated using the DSSP algorithm (Kabsch and Sander, 1983). The solvent accessible surface area was calculated for the sidechains only, using a probe radius of 1.4 Å. To compare changes in the solvent accessible surface area along the sequence necessitates correcting for the intrinsic differences in sidechain volume. To account for this, we performed simulations in which all attractive non-bonded interactions are turned off. In these simulations only repulsive component of the Lennard-Jones potential determines the energetically accessible ensemble. This excluded volume (EV) ensemble allows for the intrinsic SASA of each reside sidechain in the appropriate sequence context to be computed, as done previously (Holehouse et al., 2015). The normalized SASA is then calculated as the ratio of the per-residue SASA from the full simulation divided by the SASA from the EV ensemble. Finally, the change in normalized SASA was computed by calculating the difference between the normalized SASA in the wildtype simulation and each of the variants. Data from simulation analysis, subsampled trajectories information, and simulation input information can be found at https://github.com/holehouse-lab/supportingdata/tree/master/2022/jankowski_2022.

### AlphaFold2 structural modelling

The modeled 50 a.a. portion of FRQ (centered on the FFD region) shown in Figure 7A was done using a Google Colab notebook based on a simplified version of AlphaFold v2.1.0 (Jumper et al., 2021), made available at the following website: https://colab.research.google.com/github/deepmind/alphafold/blob/main/notebooks/AlphaFold.ipynb.

