## Supplemental Figures for "The formation of a fuzzy complex in the negative arm regulates the robustness of the circadian clock"

**Supplemental Figures and Movie**

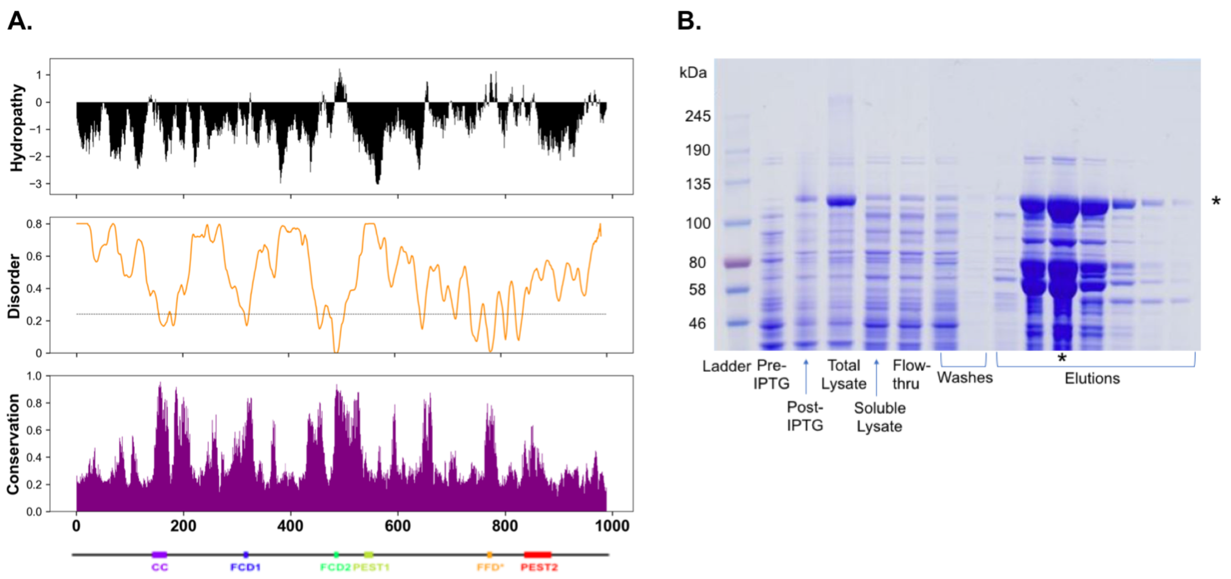


Supplemental Figure 1. FRQ is highly disordered and not amenable to typical biophysical methods, necessitating the LOCATE analysis. (A) Linear sequence analysis of FRQ. Top: Linear hydropathy profile reveals per-residue hydrophobicity based on the Kyte-Doolittle scale (Kyte and Doolittle, 1982). The majority of the sequence is highly depleted for hydrophobic residues. Middle: Linear disorder profile calculated using metrapredict, using with a five-residue smoothing window. FRQ is predicted to be almost entirely disordered. Bottom: Per-residue conservation calculated across 83 FRQ orthologs taken from eggNOG. Regions of high conservation coincide with hydrophobic regions with lower disorder tendencies. (B) A Coomassie-stained gel showing all steps of the FRHΔ100 induction (pre- and post-IPTG addition), as well as total versus soluble lysate after French press and nickel column purification steps (flow-thru, washes and final elutions). The asterisk (*) denotes the elution used in subsequent microarray assays as well as the band that corresponds to FRHΔ100. Related to Figure 1.

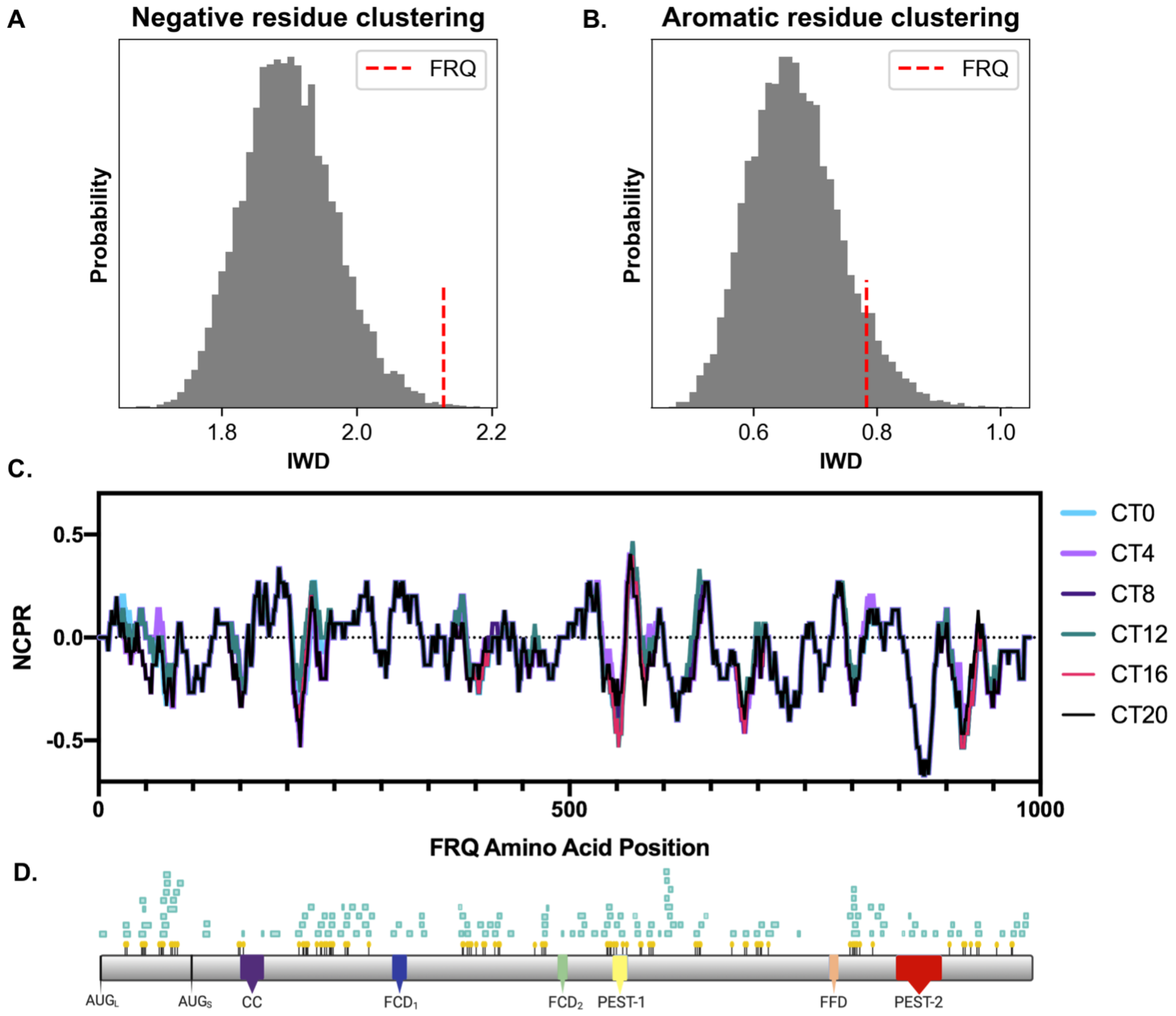


Supplemental Figure 2. FRQ sequence features imply functionally important factors for FRQ function. (A) The statistical significance of negatively charged clusters (islands) of residues is assessed by calculating the expected null distribution for negatively charged residue clustering from randomly shuffled of sequences that match the composition and length of FRQ. Clustering is calculated using the Inverse Weighted Distance (IWD) (see methods). The real FRQ sequence (red dashed line) has negatively charged residues that are more well clustered than almost all randomly shuffled sequences, suggesting that at least for FRQ from *N. cassa,* negatively residues (like positively charged residues) are well-clustered. (B) The statistical significance of aromatic clusters of residues is assessed by calculating the expected null distribution for aromatic residue clustering from randomly shuffled of sequences that match the composition and length of FRQ. Clustering is calculated using the Inverse Weighted Distance (IWD) (see methods). The real FRQ sequence (red dashed line) has aromatic residues that are not more clustered than almost all randomly shuffled sequences. (C) Net Charge Per Residue (NCPR) plot of FRQ incorporating extra negative charges at known phosphorylated residues at the different Circadian Times (CT), as published in (Baker et al., 2009). Note that CT = 0 is relative dawn and CT = 12 is relative dusk. (D) Predicted SLiMs (green boxes) from the ELM database for verified interactors of FRQ, along with detected phosphosites denoted with yellow pins (Baker et al., 2009; Gouw et al., 2017; Pelham et al., 2021). See Figure 1 for more details about the known FRQ domains highlighted in color. Related to Figure 3.

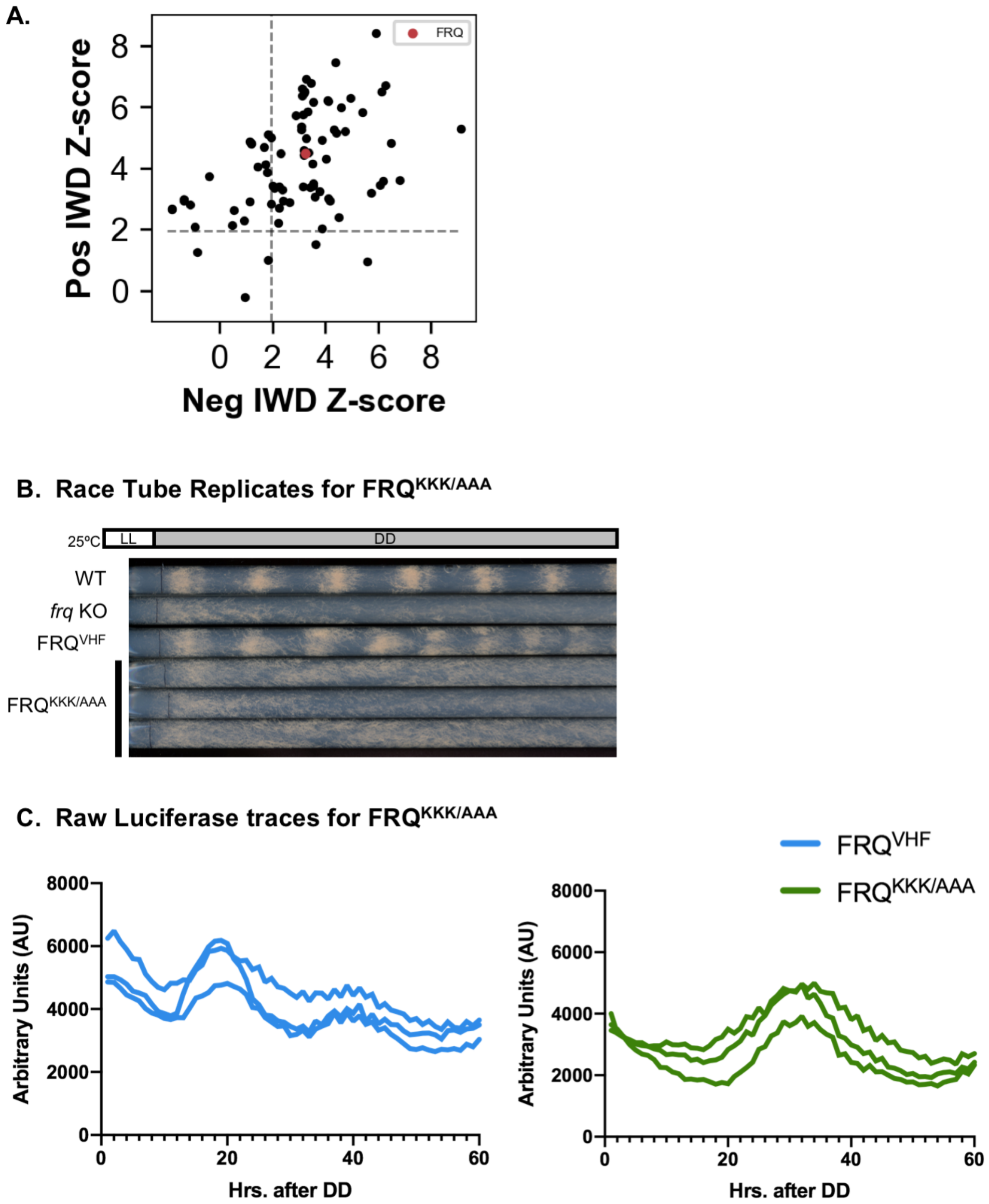


Supplemental Figure 3. Positively charged islands are a conserved across FRQ orthologs. (A) The Z-score of positive residue Inverse Weighted Distance (IWD) versus negative residue IWD of FRQ (red circle) compared to 86 FRQ orthologs (black circles). Dotted lines denote significance ($p < 0.05$). Most sequences (64/86) show significant clustering of both positively-charged and negatively-charged residues (top right quadrant). A subset of sequences (17/86) only show clustering of positively charged residues (top left quadrant). A much smaller subset show either non-significant clustering (3/86) or only clustering of negatively charged residues (2/86). (B) Race tube replicates of the FRQ$^{KKK/AAA}$ mutant strain. (C) Raw luciferase traces (N=3) from the FRQ$^{KKK/AAA}$ mutant strain 96-well plate experiment. Related to Figure 3.

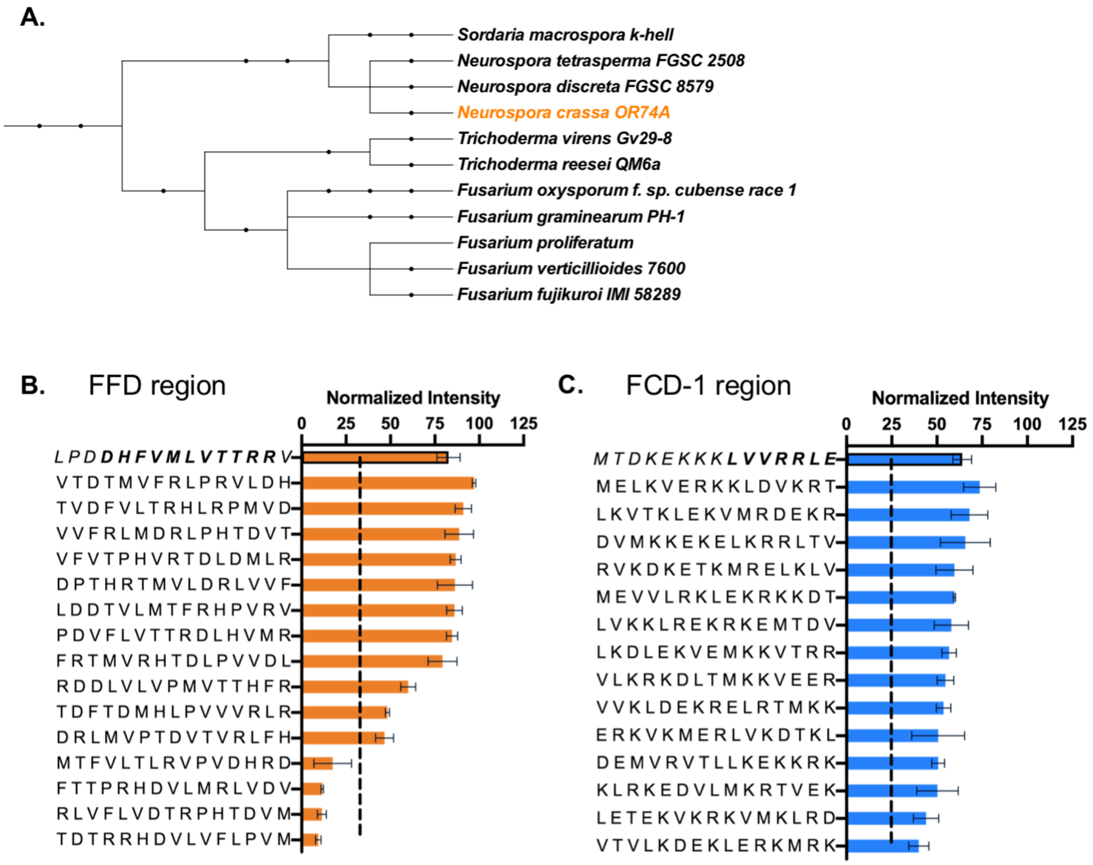


<u>Supplemental Figure 4. Sequence composition contributes to the interaction between the FRQ-FFD and FRH.</u> (A) A phylogenetic tree of the FRQ homologs considered in the FFD region alignment. (B) Normalized binding intensity of the native and scrambled peptides of the FRQ-FFD region to FRH, based on peptides from Library I, using ~10 nM FRH and visualized using anti-His. Peptide sequences are reported along the y-axis with the native FFD in bold. Normalized binding intensity values are reported for each peptide along the x-axis and error bars report the standard deviation (N=3). The dashed line represents one Standard Deviation above background. (C) Normalized binding intensity of the native and scrambled peptides of the FRQ-FCD-1 region to FRH, based on peptides from Library II, using ~100 nM FRH and visualized using anti-His. Peptide sequences are reported along the y-axis with the native FCD-1 in bold. Binding intensity values are reported for each peptide along the x-axis, and error bars report the standard deviation (N=3). The dashed line represents one standard deviation above background-subtracted zero, for the whole library. Related to Figure 4.

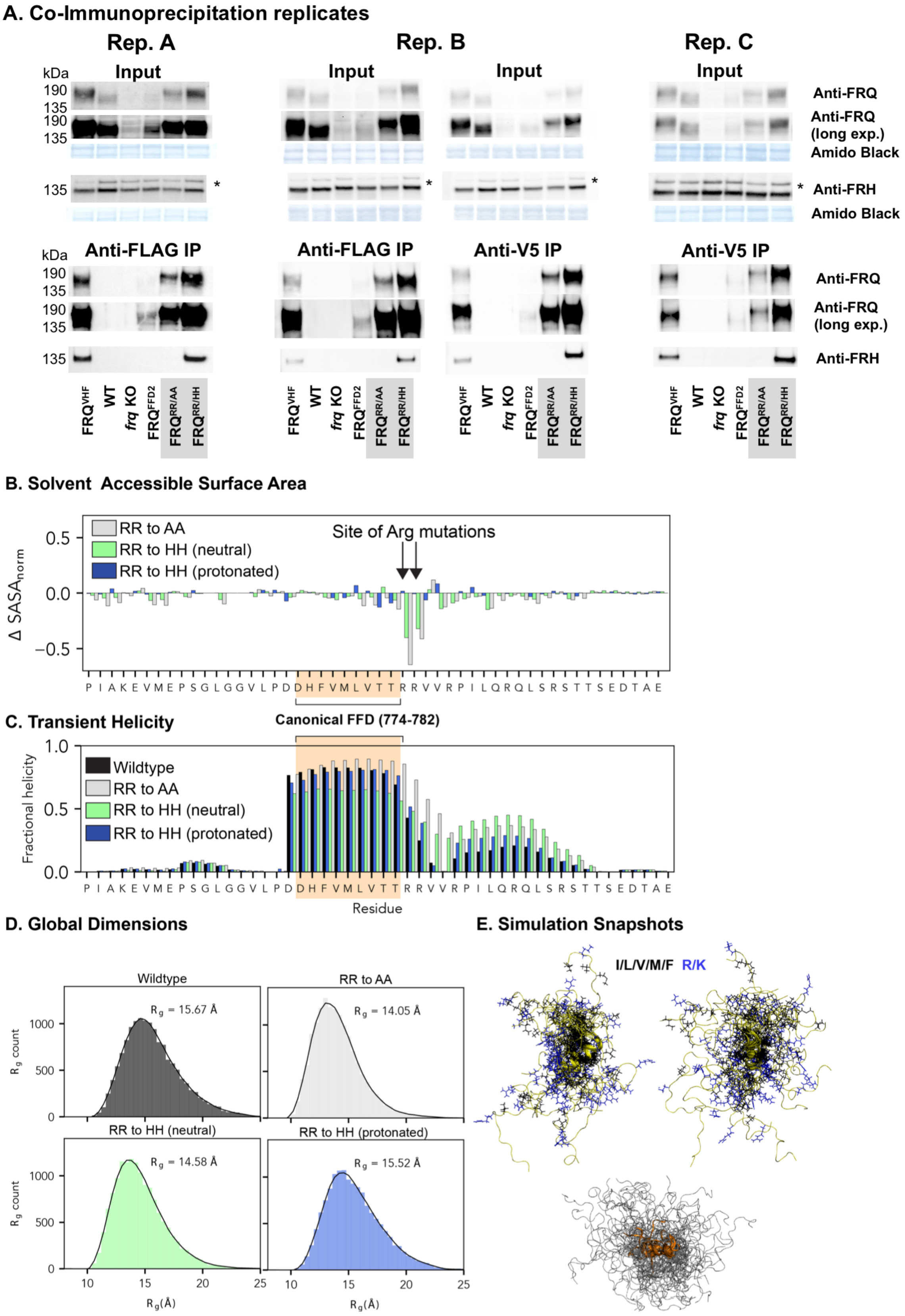
A. Co-Immunoprecipitation replicates
Rep. A
Rep. B
Rep. C
kDa
Input
190
135
Anti-FRQ
Anti-FRQ (long exp.)
Amido Black
Anti-FRH
*
Anti-FLAG IP
Anti-V5 IP
FRQ$^{VHF}$
WT
frq KO
FRQ$^{FFD2}$
FRQ$^{RR/AA}$
FRQ$^{RR/HH}$
B. Solvent Accessible Surface Area
RR to AA
RR to HH (neutral)
RR to HH (protonated)
Site of Arg mutations
$\Delta$ SASA$_{norm}$
0.5
0.0
−0.5
P I A K E V M E P S G L G G V L P D D H F V M L V T T R R V V R P I L Q R Q L S R S T T S E D T A E
Canonical FFD (774-782)
C. Transient Helicity
Wildtype
RR to AA
RR to HH (neutral)
RR to HH (protonated)
Fractional helicity
1.0
0.5
0.0
Residue
D. Global Dimensions
Wildtype
RR to AA
RR to HH (neutral)
RR to HH (protonated)
$R_g$ = 15.67 Å
$R_g$ = 14.05 Å
$R_g$ = 14.58 Å
$R_g$ = 15.52 Å
$R_g$ count
1000
500
0
10
15
20
25
$R_g$(Å)
E. Simulation Snapshots
I/L/V/M/F R/K

Supplemental Figure 5. Loss of arginine hotspot residues identified by LOCATE leads to no major change in the FFD predicted ensemble, a decrease in FRH interaction, and a decline in FRQ half-life. (A) Replicates of the Input and anti-FLAG immunoprecipitation (IP) or anti-V5 IP for the noted strains. Asterisk (*) denotes a non-specific band in the lysate lanes when using anti-FRH. (B) The change in normalized solvent accessible surface area for each residue in a 50-residue FFD region assessed by all-atom Monte Carlo simulations (see methods). Other than the arginine residues themselves, no significant changes in solvent accessibility are observed upon changing the arginine hotspot residues to alanine, neutral histidine, or protonated histidine. The canonical FFD region is highlight in orange. (C) Transient helicity reveals the canonical FFD lies along a region that is predominantly in a transient helix, a common binding mode for intrinsically disordered regions. RR783 mutations do not alter helicity profiles, with the exception of RR783AA which becomes slightly more helical. (D) Global dimensions of the four ensembles are assessed based on histograms of the radius of gyration ($R_g$), a measure of overall ensemble size. All four sequences have almost identical global dimensions. (E) Snapshots of superimposed structures from a subset of the simulations. Positively charged residues are shown in blue, hydrophobic residues in black, and the chain backbone in yellow. On the bottom, helical conformations are shown in orange, illustrating the transient and highly disordered nature of the ensemble. Related to Figure 5.

### A. Cycloheximide assay replicates

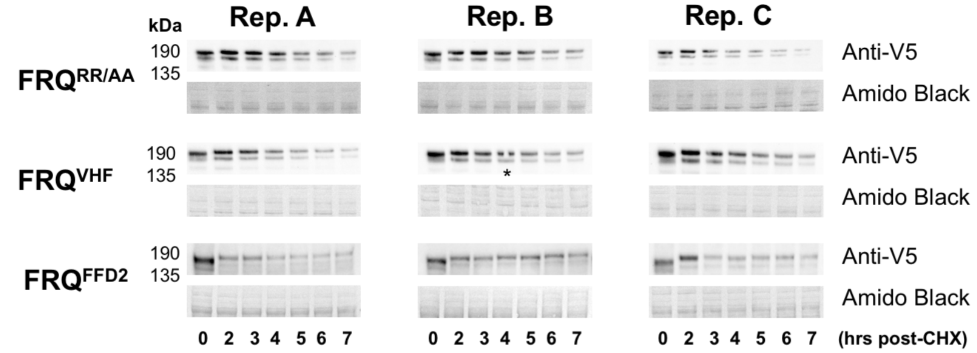


### B. Anti-WC1 lysate replicates

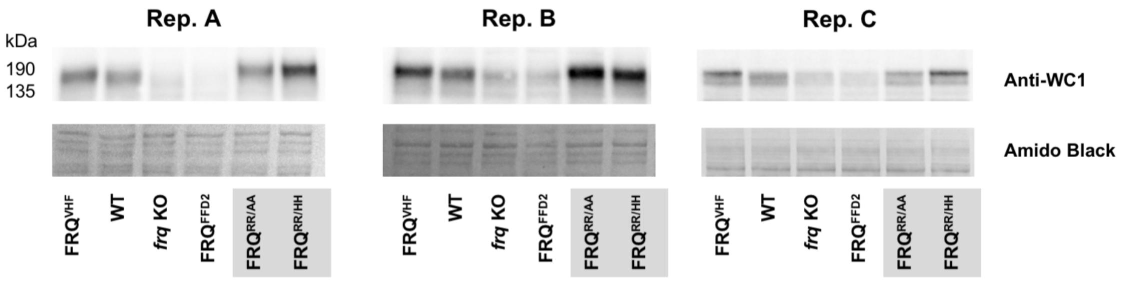


### C. Raw Luciferase Traces

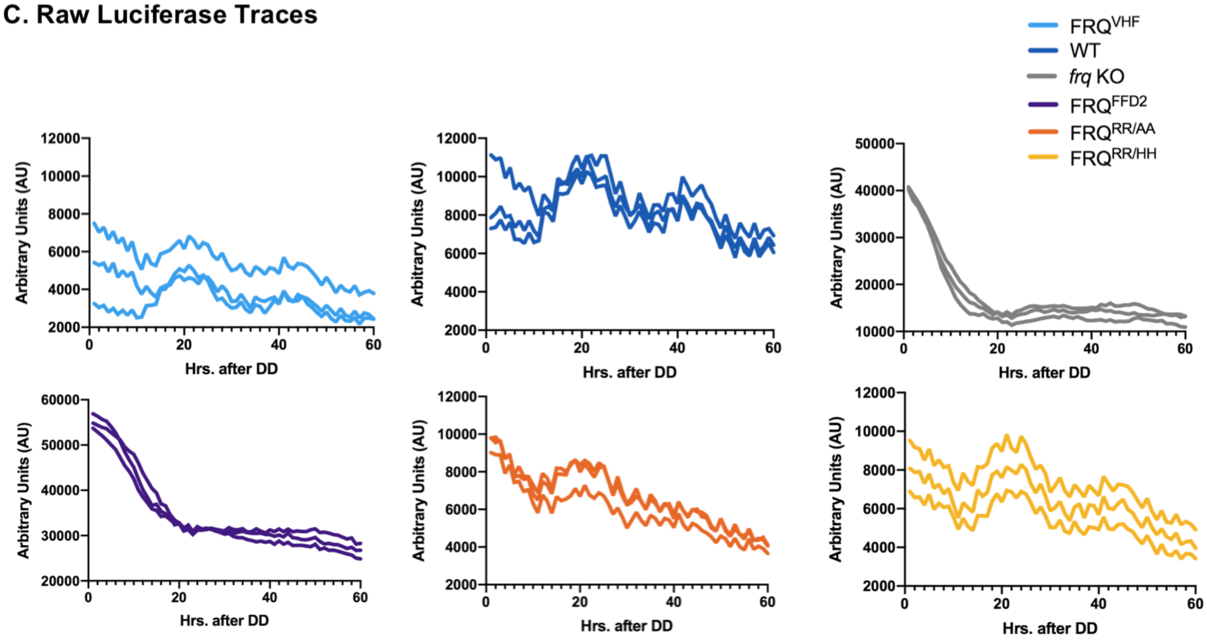

Supplemental Figure 6. FRQ$^{RR/AA}$ is more stable and supports wild-type levels of WC-1, suggesting FRQ$^{RR/AA}$ can close the TTFL independently of FRH binding. (A) Replicate blots and Amido Black stained membranes for the cycloheximide assay results presented in Figure 5C. Asterisk (*) notes that this lane was omitted from analysis due to an air bubble. (B) Replicates of anti-WC-1 lysate levels amongst the different strains, and matching Amido Black stained membranes showing even loading. Note that replicates A and B were run on Tris-Acetate gels (3-8%) while replicate C was run on a Bis-Tris gel (4-12%). (C) Raw Luciferase Traces (N=3) for each of the indicated strains, measured in Arbitrary Units (AU) over 15 min., repeated every 1 hr for 60 hrs total. Related to Figures 5 and 6.

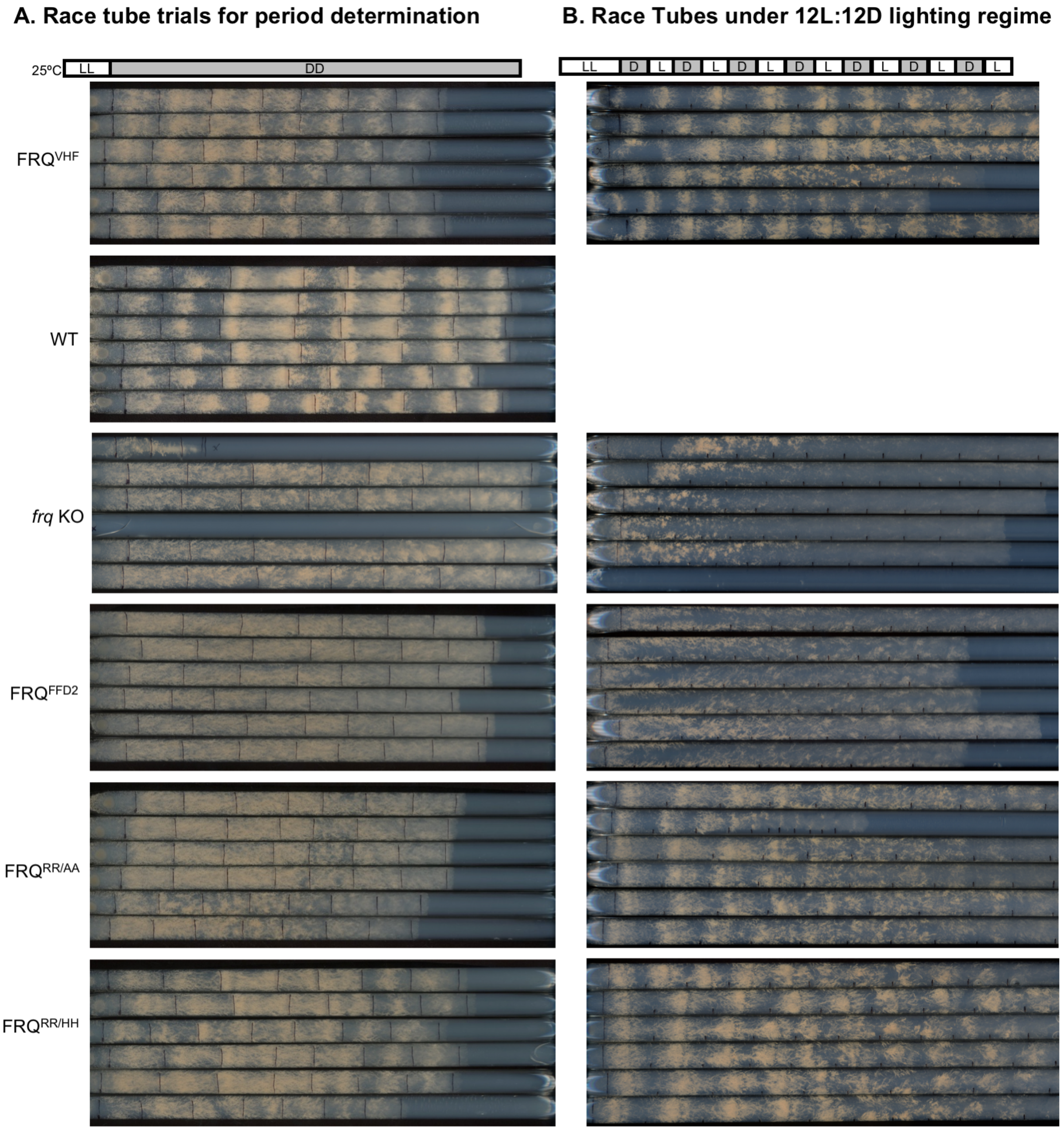


Supplemental Figure 7. FRQ$^{RR/AA}$ is overtly arrhythmic in constant conditions yet rhythmic in a 12L:12D lighting regime. (A) Race tubes grown in constant light (LL) before being allowed to free run in constant dark (DD) for each of the described *N. crassa* strains. (B) Race tubes grown in a 12 hr. light: 12 hr. dark (12L:12D) lighting regime for each of the described *N. crassa* strains. Daily marks are denoted with black lines.

Supplemental Movie 1. Movie of Monte Carlo all-atom simulation showing the overall disordered nature, albeit with transient helices, that characterizes the FFD region of FRQ. Residues 754-803 of Wild-type FRQ were simulated using the programs CAMPARI and ABSINTH (see methods for complete details). Residue and helices colored as in Supplemental Figure 5E.
